# Impaired oxidative phosphorylation drives primary tumor escape and metastasis

**DOI:** 10.1101/2025.01.08.631936

**Authors:** Emily N. Arner, Erin Q. Jennings, Daniel R. Crooks, Christopher J. Ricketts, Melissa M. Wolf, Matthew A. Cottam, Emilie L. Fisher-Gupta, Martin Lang, Nunziata Maio, Yuki Shibata, Evan S. Krystofiak, Logan M. Vlach, Zaid Hatem, Alexander M. Blatt, Darren R. Heintzman, Allison E. Sewell, Emma S. Hathaway, KayLee K. Steiner, Xiang Ye, Samuel Schaefer, Zachary A. Bacigalupa, W. Marston Linehan, Kathryn E. Beckermann, Frank M. Mason, Kamran Idrees, W. Kimryn Rathmell, Jeffrey C. Rathmell

**Affiliations:** Department of Pathology, Microbiology, and Immunology, Vanderbilt University Medical Center; Nashville, TN, USA; Department of Medicine, Vanderbilt University Medical Center; Nashville, TN, US; Urologic Oncology Branch, Center for Cancer Research, National Cancer Institute; Bethesda, MD, USA; Clinical Cancer Metabolism Facility, Center for Cancer Research, National Cancer Institute, Bethesda, MD, USA; Department of Surgery, Division of Surgical Oncology and Endocrine Surgery, Vanderbilt University Medical Center; Nashville, TN, USA; Department of Cell and Developmental Biology, Vanderbilt University; Nashville, TN, USA; Cell Imaging Shared Resource, Vanderbilt University; Nashville, TN, USA; Vanderbilt-Ingram Cancer Center, Vanderbilt University Medical Center; Nashville, TN, USA; Vanderbilt Center for Immunobiology, Vanderbilt University Medical Center; Nashville, TN, USA

**Author notes:** National Institute of Diabetes and Digestive and Kidney Diseases, National Institutes of Health; Bethesda, MD, USA.

**Keywords:** OxPhos, glycolysis, mitochondria, EMT, NDUFA4L2, Hypoxia, metastasis

## Abstract

Metastasis causes most cancer deaths and reflects transitions from primary tumor escape to seeding and growth at metastatic sites. Epithelial-to-mesenchymal transition (EMT) is important early in metastasis to enable cancer cells to detach from neighboring cells, become migratory, and escape the primary tumor. While different phases of metastasis expose cells to variable nutrient environments and demands, the metabolic requirements and plasticity of each step are uncertain. Here we show that EMT and primary tumor escape are stimulated by disrupted oxidative metabolism. Using Renal Cell Carcinoma (RCC) patient samples, we identified the mitochondrial electron transport inhibitor NDUFA4L2 as upregulated in cells undergoing EMT. Deletion of NDUFA4L2 enhanced oxidative metabolism and prevented EMT and metastasis while NDUFA4L2 overexpression enhanced these processes. Mechanistically, NDUFA4L2 suppressed oxidative phosphorylation and caused citric acid cycle intermediates to accumulate, which modified chromatin accessibility of EMT-related loci to drive primary tumor escape. The effect of impaired mitochondrial metabolism to drive EMT appeared general, as renal cell carcinoma patient tumors driven by fumarate hydratase mutations with disrupted oxidative phosphorylation were highly metastatic and also had robust EMT. These findings highlight the importance of dynamic shifts in metabolism for cell migration and metastasis, with mitochondrial impairment driving early phases of this process. Understanding mitochondrial dynamics may have important implications in both basic and translational efforts to prevent cancer deaths.

## Main Text

Escape from the primary tumor and subsequent metastasis remains the single biggest factor in cancer related deaths. The metastasis process encompasses multiple phases, including, but not limited to cancer cell escape from the primary tumor, entry to the blood stream or lymphatic vessels, extravasation to secondary sites, and colonization and expansion as metastases. Metastasis is facilitated by several transitions, most notably the acquisition of tumor cell epithelial plasticity and subsequent adoption of mesenchymal-like features by epithelial tumor cells, referred to as epithelial-to-mesenchymal transition (EMT), which enables cell migration and invasion^1^. Epithelial plasticity includes EMT and the reverse, mesenchymal-to-epithelial transition (MET), which is often required for successful metastatic colonization in the metastatic site^2–4^. Notably, while circulating tumor cells and metastatic lesions have been shown to be have increased oxidative metabolism relative to primary tumors^5,6^ additional metabolic changes would be predicted to accompany such dynamic cellular transtions. However, metabolic shifts as drivers of cell state changes or metabolic requirements of the early phases of metastasis remain poorly understood.

Tumors require an abundance of energy for malignant growth, proliferation, and metastasis and metabolic demands may change with each phase of tumorigenesis. Multiple metabolic signaling pathways have been reported to increase metastasis such as glycolysis and fatty acid metabolism^7^. Decreased one-carbon metabolism through loss of the serine biosynthetic pathway rate limiting enzyme, phosphoglycerate dehydrogenase (PHGDH), can also drive metastasis in breast cancer by increasing protein glycosylation and subsequent cell migration and invasion^8^. Elevated glycolytic metabolism was observed by metabolic tracing in patient primary clear cell renal cell carcinoma (ccRCC) tumors compared with adjacent normal kidney^9,10^. However, ccRCC metastases displayed a more oxidative metabolic profile, suggesting a plasticity in mitochondrial repiratory chain activity in ccRCC throughout disease progression^10^.

Here we identified essential metabolic features of the EMT transition state in renal cell carcinoma (RCC) metastasis. The mitochondrial electron transport chain inhibitor NDUFA4L2 was upregulated in RCC patient samples and mouse models undergoing EMT. NDUFA4L2 functioned as a metabolic switch that inhibited oxidative phosphorylation (OxPhos), promoting primary tumor escape and spontaneous metastasis from orthotopic tumors. Converesly, loss of NDUFA4L2 prevented EMT and metastasis. Mirroring these results, highly metastatic human RCC characterized by loss of fumarate hydratase (FH) also exhibited impaired OxPhos that led to robust EMT. These results show that EMT is facilitated by the inhibition or loss of mitochondrial oxidative metabolism to enable primary tumor escape.

### NDUFA4L2 is associated with EMT and predicts poor survival in renal cell carcinoma

Metabolic mechanisms governing EMT are largely unknown. We thus performed bioinformatic analysis of publicly available single-cell RNA sequencing (scRNA-seq) from RCC patient primary and metastatic tumors, 7 of which were ccRCC and 1 papillary RCC (pRCC)^11^. We identified a tumor cell population characterized by EMT genes, *SNAI2* and *VIM*, which encode Slug and Vimentin, respectively (**Extended Data** Figure 1**, A-C**). This population was found in the TP2 tumor cell population, previously characterized by metabolic and glycolytic gene signatures^11^. Further analyses identified expression of NADH dehydrogenase [ubiquinone] 1 alpha subcomplex, 4-like 2 (*NDUFA4L2*) as correlated with expression of both EMT genes *VIM* and *IGFBP3* and glycolytic genes *LDHA* and *ENO1* (**Extended Data** Figure 1D). Consistent with these results, *NDUFA4L2* has been reported as a HIF1α induced inhibitor of mitochondrial electron transport complex I^12^, and complex IV of the electron transport chain^13^. Bioinformatic analyses of an independent ccRCC patient scRNA-seq dataset^14^ identified *NDUFA4L2* as highly associated with mesenchymal genes, *TGFB1* and *CDH2*, but not the epithelial gene *CDH1*, which encodes E-Cadherin (**Fig. 1A-E**). The majority of *NDUFA4L2* expression was found in tumor cells, but not normal kidney or immune cells (**Extended Data** Figure 2A).

**Fig. 1.**
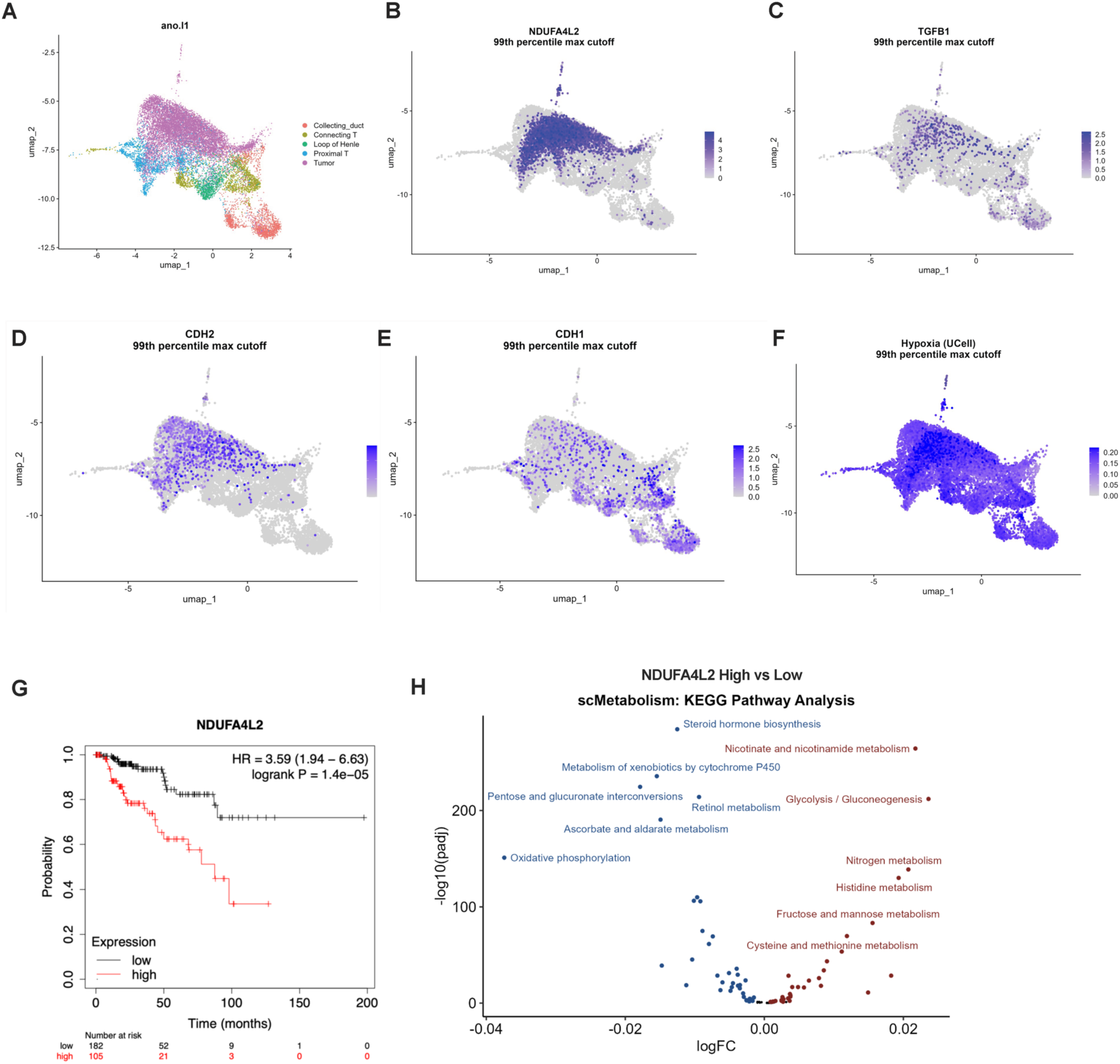
NDUFA4L2 is associated with EMT and predicts poor survival in ccRCC. **(A)** scRNAseq expression clustering from GEO accession #GSE178481^14^ for tumor cells from ccRCC evaluating the expression of **(B)** *NDUFA4L2*, **(C)** *TGFB1*, **(D)** *CDH2*, **(E)** *CDH1*, and **(F)** hallmark hypoxia signature. **(G)** Using TCGA data, high gene expression of *NDUFA4L2* in ccRCC tumors was identified as a unfavorable prognosis marker in ccRCC. High expression is in red and low expression is in black. **(H)** Using the scRNAs seq data from (A), the scMetabolism package^31^ was used to estimate pathway activity for KEGG metabolic pathways, and the Wilcoxon rank sum test was used to detect differential activity of metabolic pathways. Red pathways represent metabolic pathways upregulated in *NDUFA4L2* high cells, while the blue pathways represent metabolic pathways upregulated in *NDUFA4L2* low cells.

Data from the Cancer Genome Atlas (TCGA) database showed that *NDUFA4L2* expression was higher in ccRCC compared with normal kidney or other cancer types (**Extended Data** Figure 2B). This was ccRCC specific, as *NDUFA4L2* was minimally expressed in tumors of patients with primary papillary RCC (pRCC) (**Extended Data** Figure 2**, B and C**), although the expression levels were variable. Samples from a ccRCC patient with bone metastases also had high expression of *NDUFA4L2* in the primary tumor (**Extended Data** Figure 2C). NDUFA4L2 has been reported as a transcriptional target of HIF1α^12^, therefore it is possible that *NDUFA4L2* expression is highest in ccRCC due to widespread loss of VHL and stability of HIF1α. Indeed, *NDUFA4L2* high cells in patient scRNA-seq data had the highest expression of hypoxia signature genes (**Fig. 1F**). Survival analysis from TCGA data revealed that higher gene expression (top 50% based on median expression) of *NDUFA4L2* correlated with decreased overall survival in RCC (**Fig. 1G)**. The correlation between *Ndufa4l2* and EMT was also found in a discrete subset of cells by scRNA-seq of syngeneic RCC murine subcutaneous tumors with normal *VHL* expression^15^ (**Extended Data** Figure 2**, D and E**). To evaluate the role NDUFA4L2 in metabolism using patient RCC samples, metabolic pathway analysis in the patient scRNA-seq dataset^14^ identified that *NDUFA4L2* high expressing tumor cells had upregulated signatures of glycolysis/gluconeogenesis while *NDUFA4L2* low expressing tumor cells had upregulated OxPhos. (**Fig. 1H)**.

### NDUFA4L2 inhibits oxidative phosphorylation

RCC patient primary tumors displayed high glycolysis and lower oxidative phosphorylation rates compared with normal adjacent kidney. Extracellular flux analyses of fresh ccRCC tumor samples from two patients showed higher extracellular acidification rate (ECAR) and lower oxygen consumption rate (OCR) in the tumor cells compared with adjacent normal kidney, indicating a shift toward increased glycolysis in ccRCC tumors (**Fig 2, A and B**). To determine the impact of NDUFA4L2 on RCC tumor cell metabolic phenotype in vitro and in vivo, clonal NDUFA4L2 deficient (KO) cell lines were generated in syngeneic RENCA and *VHL*-deficient LVRCC67^16^ RCC cell lines. To establish if changes seen in the KO cells were due to loss of NDUFA4L2 expression and not off target effects, the KO cell lines were transduced to stably rescue and overexpress NDUFA4L2 expression (**Fig. 2, C and D, and Extended Data** Figure 3**, A to C**). RNA sequencing of the EV, KO, and rescue RENCA cell lines found biological oxidations, extracellular matrix organization, and metabolism pathways were highly impacted by modulation of NDUFA4L2 expression (**Extended Data** Figure 3D). These findings are consistent with the correlation of EMT and impaired oxidative phosphorylation with *NDUFA4L2* in patient samples.

**Fig. 2.**
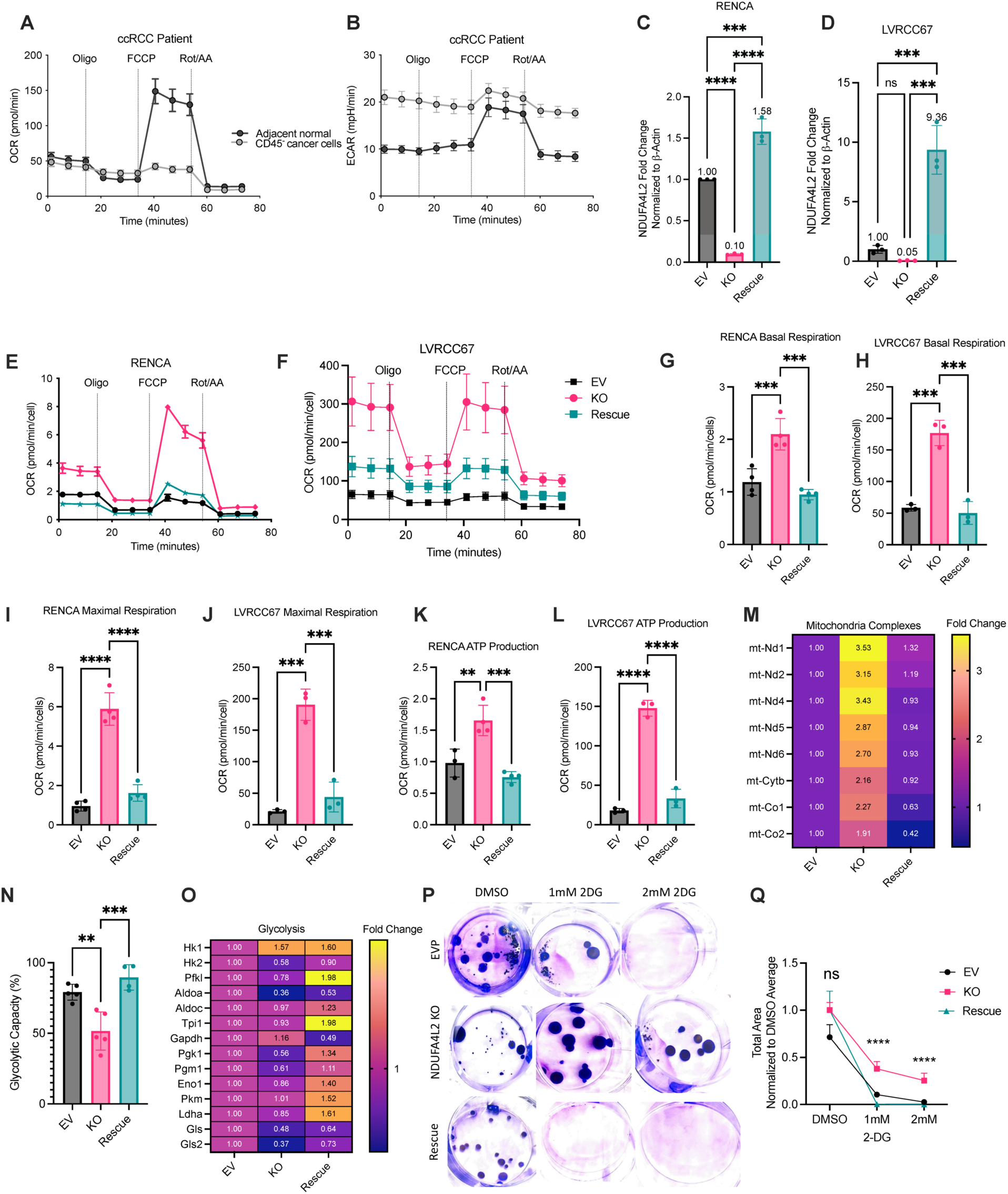
NDUFA4L2 inhibits oxidative phosphorylation. CD45 negative cells were isolated from a fresh patient ccRCC tumor and normal adjacent kidney. Extracellular flux analysis was done on the isolated cells and **(A)** oxygen consumption rate (OCR) and **(B)** extracellular acidification rate (ECAR) were measured. Results are representative of two individual patients. Gene expression of NDUFA4L2 in EV, NDUFA4L2 KO, and rescue **(C)** RENCA and **(D)** LVRCC67 cell lines. Gene expression was measured using qPCR and normalized to β-Actin. Fold change was plotted by normalizing expression to the empty vector (EV). Each dot represents an individual experiment, with 3-6 technical replicates/experiment. Extracellular flux analysis was done on EV, KO, and rescue **(E)** RENCA and **(F)** LVRCC67 cells. The graphs are representative of 3 or more individual experiments, with 3-4 biological replicates and 6 technical replicates each. OCR and ECAR measurements were normalized to total cell number in the well. Basal respiration was calculated in **(G)** RENCA and **(H)** LVRCC67 cells, maximal respiration in **(I)** RENCA and **(J)** LVRCC67 cells, and ATP production in **(K)** RENCA and **(L)** LVRCC67 EV, KO and rescue cells. Each dot represents a biological replicate. **(M)** RNA-seq of RENCA EV, KO, and rescue cells. Three technical replicates were sequenced per cell line. Gene counts were normalized to the average of the EV gene counts for each gene. **(N)** SCENITH of EV, KO, and rescue RENCA cells treated with puromycin, 2DG, oligomycin individually and in combination. Protein translation by puromycin incorporation was used to calculate glycolytic capacity (%) in each condition. Each dot represents a biological replicate combined from 2 individual experiments. **(O)** RNA-seq of RENCA EV, KO, and rescue cells revealed that glycolytic transcripts are increased in EV and rescue cells compared to NDUFA4L2 KO cells. Three technical replicates were sequenced per cell line. Gene counts were normalized to the average of the EV gene counts. **(P)** EV, KO, and rescue RENCA cells were plated at low confluence (∼50 cells/well) in a 6-well plate and treated with DMSO, 1mM 2DG or 2mM 2DG for 2 weeks. Cells were fixed and stained with crystal violet. **(Q)** Total area of crystal violet/well was quantified using Image J and normalized to the average of the DMSO wells/condition. 6 wells/condition were quantified for each experiment and the experiment was repeated 3 times. Statistics were done using a one-way anova with multiple comparisons, **p<0.01, ***p<0.001, ****p<0.0001.

Comparable to patient ccRCC cells (**Fig. 2B**), NDUFA4L2 expressing RENCA and LVRCC67 cells had lower OCR than cells lacking NDUFA4L2 (**Fig. 2, E and F, and Extended Data** Figure 4**, A and B**). Conversely, resembling the human adjacent normal samples, NDUFA4L2 KO cells had increased basal (**Fig. 2, G and H**) and maximal respiration (**Fig. 2, I and J**), ATP production (**Fig. 2, K and L**), and spare respiratory capacity (**Extended Data** Figure 4**, C and D**). NDUFA4L2 thus inhibits oxidative phosphorylation. To metabolically profile NDUFA4L2 EV, KO, and rescue RENCA cells at a single-cell resolution, we utilized a single cell-based assay for puromycin incorporation as an indicator of protein synthesis rates and therefore metabolism, referred to as SCENITH^17^. Using this method, NDUFA4L2 KO cells had higher mitochondrial dependence compared with EV or rescue cells (**Extended Data** Figure 4E). Interestingly, bulk RNA-sequencing identified multiple mitochondrial encoded electron transport chain transcripts elevated in NDUFA4L2 KO cells compared with EV or rescue, suggesting NDUFA4L2 may inhibit mitochondrial transcription or suppress mitochondria abundance (**Fig. 2M**).

Glycolytic rates were reciprocally regulated with OxPhos. ECAR and glycolytic capacity were reduced as OCR increased in NDUFA4L2 KO cells compared with EV and rescue cells (**Fig. 2N, and Extended Data** Figure 4**, F and G**). RNA-sequencing and qRT-PCR identified select glycolytic transcripts as decreased in NDUFA4L2 KO cells compared with EV or rescue (**Fig. 2O, and Extended Data** Figure 4**, H and I**). Consistent with reduced dependence on glycolysis, NDUFA4L2 KO cells were more resistant to glycolysis inhibition upon culture with 2-deoxyglucose (2DG) than NDUFA4L2 EV or rescue cells, which each showed high levels of cell death with 2DG (**Fig. 2, P and Q**). Given the striking inhibition of OxPhos by NDUFA4L2, we visually profiled mitochondria of EV, KO, and rescue cells using transmission electron microscopy (TEM). Cells with intact NDUFA4L2 had mitochondria with few cristae, partially explaining why cells with NDUFA4L2 have very little oxidative phosphorylation activity and heavy reliance on glycolysis (**Extended Data** Figure 5A). Interestingly, cell morphological differences were also notable in the TEM images, as EV and rescue cells had a more elongated and mesenchymal shape while KO cells appeared more rounded (**Extended Data** Figure 5A). Although cellular reactive oxygen species (ROS) and mitochondrial ROS levels were not significantly changed (**Extended Data** Figure 5B and C), mitochondrial mass (Mitotracker) and mitochondrial membrane potential (TMRE) were increased in NDUFA4L2 KO cells compared with EV (**Extended Data** Figure 5**, D and E**). Multiple mitochondrial regulatory genes were affected by NDUFA4L2 (**Extended Data** Figure 5F). NDUFA4L2 expression thus impairs oxidative phosphorylation and increases reliance on glycolysis.

### NDUFA4L2 regulates chromatin accessibility of EMT genes

Given the effect of NDUFA4L2 on metabolism and EMT in RCC, we hypothesized the metabolic effects of NDUFA4L2 inhibition on oxidative phosphorylation may drive EMT gene expression. To better understand the metabolic function of NDUFA4L2, targeted metabolomics were performed on EV, NDUFA4L2 KO, and rescue RENCA cells. Surprisingly, intracellular glucose was increased in NDUFA4L2 KO cells compared with EV or rescue (**Extended Data** Figure 6A), although downstream metabolites levels remained mostly unchanged (**Extended Data** Figure 6**, B to I**). Two metabolites that were notably differentially regulated by NDUFA4L2 expression were α-Ketoglutarate (α-KG) and succinate, for which NDUFA4L2 expression led to elevated abundance of α-KG and blunted succinate abundance compared to NDUFA4L2 KO cells (**Fig. 3, A and B**). Chromatin-modifying enzymes often use metabolic intermediates as cofactors to regulate histone methylation^18,19^, including α-KG, which is a required cofactor for histone demethylases and regulates chromatin accessibility; while succinate can inhibit demethylation of histones via the inhibition of histone lysine demethylases (KDMs)^20^. We therefore tested if NDUFA4L2 altered histone methylation and chromatin accessibility. Using liquid chromatography and tandem mass spectroscopy (LC-MS/MS) to assess methylation status, we observed that cells with higher expression of NDUFA4L2 (EV and rescue cells) had lower levels of methylation on both cytosolic and chromatin arginine residues while total, unmodified protein arginine levels were similar in both the cytosolic and chromatin fractions (**Fig. 3, C to F, Extended Data** Figure 6**, J to M**). These data suggest NDUFA4L2 expression results in increased arginine histone demethylation, possibly via α-KG and succinate.

**Fig. 3.**
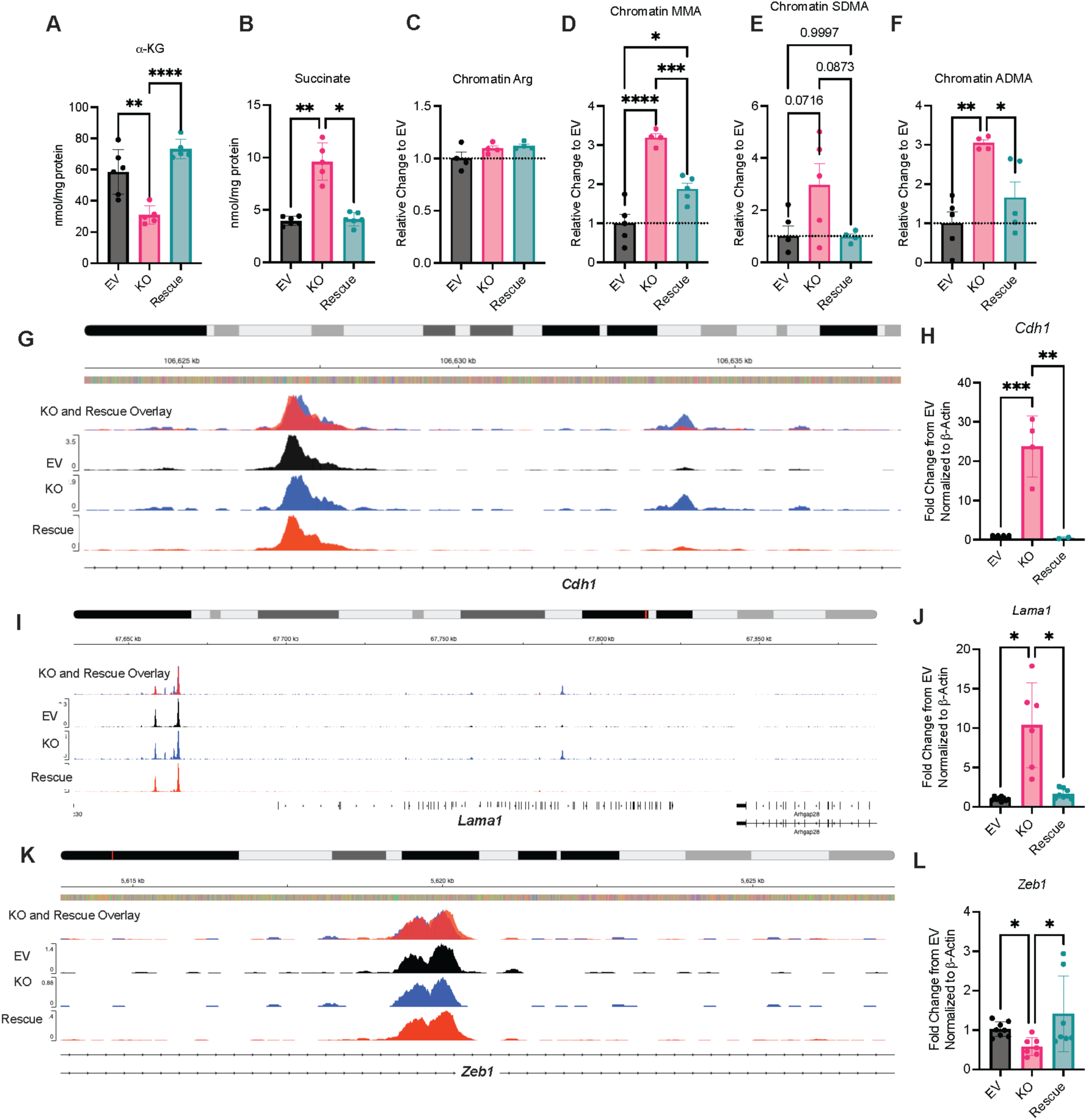
NDUFA4L2 metabolically regulates chromatin accessibility of EMT genes. LC-MS/MS was used to perform targeted metabolomics on EV, KO, and rescue RENCA cells. **(A)** α-KG and **(B)** succinate were quantified using a ^13^C-1-Lactate internal standard and normalized to the protein in each respective sample’s cell pellet. Relative change was calculated by normalizing to the average of the EV cells. N=5-6 replicates and is representative from 2 individual experiments. Quantification of arginine post-translational modifications in the chromatin of EV, NDUFA4L2 KO, and rescue RENCA cells. Quantification of **(C)** total unmodified arginine, **(D)** monomethylarginine (MMA), **(E)** symmetric dimethylarginine (SDMA), and **(F)** asymmetric dimethylarginine (ADMA). Relative change was calculated by normalizing to the average of the EV cells. N= 4-5 replicates and is representative from 2 individual experiments. Locus accessbility plots for **(G)** *Cdh1.* Overlay is of NDUFA4L2 KO and Rescue. Black represents EV, blue represents NDUFA4L2 KO, and red represents rescue. 2-3 biological replicates were sequenced per group. **(H)** Quantification of *Cdh1* gene expression normalized to β-Actin by qPCR in RENCA EV, NDUFA4L2 KO, and rescue cells. Fold change was plotted by normalizing expression to the empty vector (EV). Each dot represents an individual experiment, with 3-6 technical replicates/experiment. **(I)** Locus accessbility plots for *Lama1.* Overlay is of NDUFA4L2 KO and Rescue. Black represents EV, blue represents NDUFA4L2 KO, and red represents rescue. 2-3 biological replicates were sequenced per group. **(J)** Quantification of *Lama1* gene expression normalized to β-Actin by qPCR in RENCA EV, NDUFA4L2 KO, and rescue cells. Fold change was plotted by normalizing expression to the empty vector (EV). Each dot represents an individual experiment, with 3-6 technical replicates/experiment. **(K)** Locus accessibility plots for *Zeb1*. Overlay is of NDUFA4L2 KO and Rescue. Black represents EV, blue represents NDUFA4L2 KO, and red represents rescue. 2-3 biological replicates were sequenced per group. **(L)** *Zeb1* gene expression was measured using qPCR in RENCA EV, NDUFA4L2 KO, and rescue cells and normalized to β-Actin. Fold change was plotted by normalizing expression to the empty vector (EV). Each dot represents an individual experiment, with 3-6 technical replicates/experiment. Statistics were done using a one-way anova with multiple comparisons, *p<0.05, **p<0.01, ***p<0.001, ****p<0.0001.

As chromatin methylation is significantly altered by NDUFA4L2, we next evaluated chromatin accessibility of EMT genes using ATAC-sequencing and identified multiple EMT-associated loci in which NDUFA4L2 modulation affected chromatin accessibility. Epithelial genes, including *Cdh1* and the laminin gene *Lama1* had more open chromatin accessibility in the NDUFA4L2 KO cells compared to EV or rescue, which correlated with increased levels of these genes in the KO cells (**Fig. 3, G to J, Extended Data** Figure 7**, A and B)**. Conversely, the mesenchymal gene *Zeb1* also showed regions with altered accessibility (**Fig. 3K, Extended Data** Figure 7C), and decreased expression with NDUFA4L2 deficiency (**Fig. 3L**). These results suggest that metabolic changes in α-KG and succinate in response to NDUFA4L2 regulate arginine histone methylation and chromatin accessibility to drive EMT and primary tumor escape.

### NDUFA4L2 drives EMT and primary tumor escape

Because NDUFA4L2 regulated the chromatin accessibility of EMT-related genes, we evaluated the EMT capabilities of RCC cell lines with and without NDUFA4L2. The epithelial marker *CDH1* (ECAD) mRNA and protein expression were significantly increased in NDUFA4L2-deficient cells compared with EV and rescue cells (**Fig 3H**, **Fig. 4, A to C**). Conversely, the mesenchymal marker *CDH2* (NCAD) was decreased in NDUFA4L2 KO RENCA cells suggesting NDUFA4L2 promotes the loss of adhesion for cells to undergo EMT (**Fig. 4D**). To evaluate the functional ability of NDUFA4L2 to promote invasion in an extraceullar matrix (ECM), cells were plated in a mixture of collagen and matrigel and allowed to grow over 4 days. In the absence of NDUFA4L2, the cell colonies remained round and epithelial-like. However, when NDUFA4L2 was re-expressed, cells became invasive throughout the ECM, characteristic of mesenchymal cells (**Fig. 4E**). To evaluate if NDUFA4L2 influences EMT in vivo, EV and KO cells were injected subcutaneously into immunocompetent mice and tumors were evaluated after 3 weeks. Surprisingly, primary tumors were not significantly different in size, despite altered tumor cell metabolism (**Extended Data** Figure 8A). However, tumors without NDUFA4L2 appeared more epithelial, with higher ECAD expression and lower SLUG expression compared with EV tumors (**Fig. 4F**). NDUFA4L2 thus promotes mesenchymal phenotypes in vivo as well as in vitro.

**Fig. 4.**
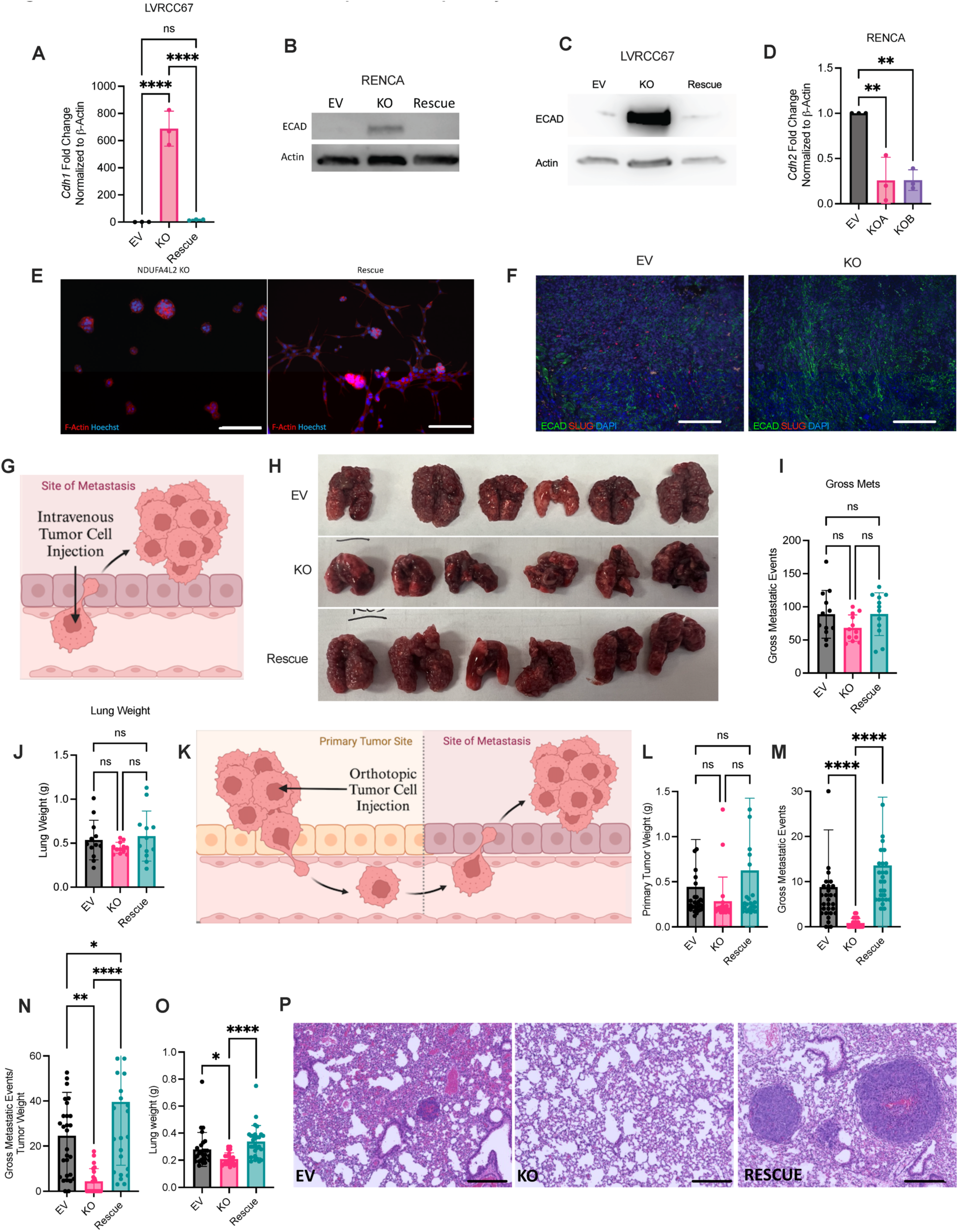
NDUFA4L2 drives EMT and escape from the primary tumor. *Cdh1* gene expression was measured using qPCR in **(A)** LVRCC67 EV, NDUFA4L2 KO, and rescue cells and normalized to β-Actin. Fold Change was plotted by normalizing expression to the empty vector (EV). Each dot represents an individual experiment, with 3-6 technical replicates/experiment. Western blot analysis of ECAD protein expression in 40 µg of protein lysed from EV, KO, and rescue **(B)** RENCA and **(C)** LVRCC67 cell lines. Each blot is representative from at least 3 individual experiments for each cell line. **(D)** *Cdh2* gene expression was measured using qPCR in EV RENCA and two different NDUFA4L2 KO clonal cell lines. mRNA was normalized to β-Actin. Fold Change was plotted by normalizing expression to the empty vector (EV). Each dot represents an individual experiment, with 3-6 technical replicates/experiment. **(E)** Representative image of NDUFA4L2 KO cells and rescue RENCA cells plated in a mixture of 70% Collagen/30% matrigel. After 4 days, cells were stained for F-Actin (red) and Hoechst (blue) and imaged using Z-stacks and confocal microscopy. Scale bars are 100 µm. **(F)** Representative images of EV, KO, and rescue subcutaneous tumors stained for ECAD (green) and SLUG (red). Scale bars are 100 µm. **(G)** Schematic of IV retro-orbital tumor cell injection in EV, KO, and rescue RENCA cells in lung colonization model. **(H)** Representative images of lungs from lung colonization. **(I)** Quantification of gross metastatic events. **(J)** Lung weight was measured in grams. Graphs show n=11-12 mice/group combined from 2 independent experiments. Two additional experiments were completed with similar results. **(K)** Schematic of EV, NDUFA4L2 KO, and rescue cells orthotopically implanted into the subrenal capsule of immunocompetent mice. **(L)** Primary tumor weight (g) measured three weeks post tumor implantation. **(M)** Quantification of gross metastatic events in the lungs. **(N)** The number of gross metastatic events were normalized to tumor weight for each mouse. **(O)** Lung weight for each mouse was measured in grams. **(P)** Representative 10X images from the lung of mice injected orthotopically with EV, KO, and rescue RENCA cells. N=23-28 mice/group combined from 5 independent experiments. Statistics were done using a one-way anova with multiple comparisons, ns = not significant, *p<0.05, **p<0.01, ***p<0.001, ****p<0.0001.

Because metastasis involves proximal steps for EMT to escape from the primary tumor and distal steps that allow seeding and growth at the metastatic niches, we tested if NDUFA4L2 affected tumor escape or growth at metastatic sites. To test if NDUFA4L2 regulated seeding metastatic niches, tumor cells were first injected directly into the bloodstream to bypass the need to tumor escape (**Fig. 4G**). There were, however, no significant differences in gross metastases or lung weight between EV, KO, or rescue (**Fig. 4, H to J**). Lung seeding and colonization do not appear affected by NDUFA4L2. To test if NDUFA4L2-induced EMT promotes escape from the primary tumor and subsequent spontaneous metastasis, EV, KO, and rescue cells were injected orthotopically into the sub renal capsule of immunocompetent mice (**Fig. 4K**). Spontaneous metastases to the lung were then evaluated. Although primary tumor weights did not differ (**Fig. 4L**), mice that received NDUFA4L2 KO cells had significantly fewer metastases than those injected with NDUFA4L2-expressing cells (**Fig. 4M, N**). Total lung weights were also decreased in mice injected with NDUFA4L2 KO cells, showing lower overall tumor burden in NDUFA4L2 KO cell lungs (**Fig. 4, O and P**). Notably, the metastatic burden correlated with NDUFA4L2 expression level, with rescue cells with the highest expression of NDUFA4L2 having the highest metastatic burden (**Fig. 2C**). Importantly, NDUFA4L2 KO cells in orthotopic primary tumors had decreased levels of glycolytic transcripts compared to cells with intact NDUFA4L2 (**Extended Data** Figure 8**, B to D**). These results highlight that NDUFA4L2 supports EMT for primary tumor escape, but does not affect metastatic site colonization.

### Decreased oxidative phosphorylation drives EMT and metastasis

To test if NDUFA4L2 regulation of EMT is specific or due more broadly to inhibition of OxPhos, we next tested alternative means to modulate mitochondrial metabolism. Mitochondrial complex I can be replaced with the yeast protein, NDI1, to override blocks on electron transport^21^. Interestingly, increasing mitochondrial electron flow in wildtype cells with NDI1 expression alone was sufficient to increase the expression of the epithelial marker ECAD (**Fig. 5A**), showing that increased OxPhos can stabilize epithelial states.

**Fig. 5.**
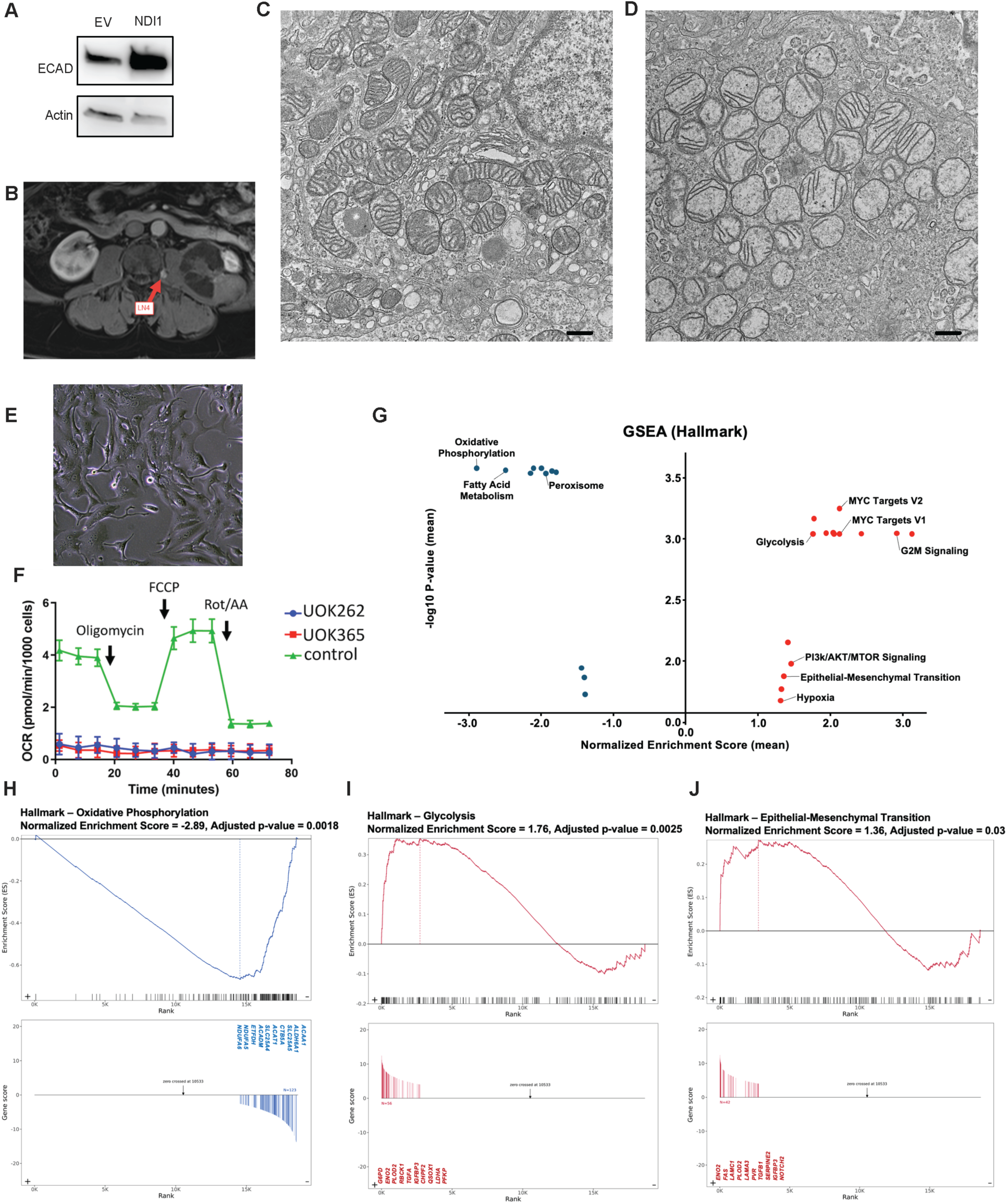
Decreased oxidative phosphorylation drives EMT and metastasis. **(A)** Western blot analysis of the yeast complex I protein NDI1 stable expression in wild-type RENCA cells. Western blot analysis of ECAD protein expression in 40 µg of protein lysed from EV and NDI1 expressing RENCA cells. **(B)** Abdominal MRI demonstrating a peri-renal lymph node metastasis in a 70-year-old male HLRCC patient. Mitochondria imaged using Transmission Electron Microscopy (TEM) in **(C)** normal adjacent kidney and **(D)** tumor from the HLRCC patient in (B). Scale bars are 0.5 µm. **(E)** Tumor cell culture from patient in (B), designated as the continuous tumor cell line UOK365. **(F)** Seahorse flux analyses of FH^-/-^ UOK262 and UOK365 tumor cells and FH^+/+^ control cells. **(G)** GSEA Hallmark analysis of bulk-RNA sequencing of 17 primary HLRCC tumors, 12 HLRCC metastatic tumors, 6 HLRCC normal adjacent kidneys, and 12 normal kidneys from donors. While **(H)** oxidative phosphorylation was enriched in normal kidney compared with HLRCC tumor, **(I)** glycolysis and **(J)** EMT were enriched in tumors compared with normal kidneys.

We next examined EMT in tumors with known mitochondrial deficiencies. Hereditary leiomyomatosis and renal cell cancer (HLRCC), is a rare and extremely metastatic RCC caused by inactivating mutations in the fumarate hydratase (*FH*) gene^22^. These mutations disrupt the tricarboxylic acid (TCA) cycle and irreversibly uncouple oxidative respiration, resulting in increased glycolysis through loss and mutation of tumor cell mtDNA^23^. To evaluate the metabolism and EMT of this form of RCC we first evaluated the mitochondria from a 70-year-old male HLRCC patient with peri-renal lymph node metastasis (**Fig. 5B, and Extended Data** Figure 9A). Compared to mitochondria from normal adjacent kidney (**Fig. 5C**), mitochondria from the HLRCC tumor were poorly shaped and had poor cristae formation (**Fig. 5D**), mirroring murine RCC cancer cells with intact NDUFA4L2 (**Extended Data** Figure 5A). A stable cell line was generated from this tumor, UOK365^24^, which had a spindle and “EMT-like” morphology (**Fig. 5E**). *FH*^-/-^ UOK365 cells were analyzed using both next generation and Sanger mtDNA sequencing and found to harbor a homoplasmic 19 base pair deletion in the mitochondrial encoded MT-ND5 gene, resulting in a frameshift mutation (p.Phe478fs) that truncates the MT-ND5 protein product by 124 residues, or approximately 21% of the total protein length. The *MT-ND5* mutation was also present in DNA obtained from the patient’s tumor at a heteroplasmy level of 73.4%, but was not present in the germline blood DNA (**Extended Data** Figure 9B). Native immunoblot evaluation of UOK365 as well as additional tumor cell lines derived from *FH*-deficient tumors^23,25^, identified decreased mitochondrial electron complex I and complex IV compared with RPTEC control cells (**Extended Data** Figure 9**, C and D**). Extracellular flux analysis of two *FH*-deficient patient cell lines confirmed the cell lines exhibited nearly undetectable mitochondrial respiration compared to the control cells with intact *FH* (**Fig. 5F**).

Bulk-RNA sequencing was performed on 17 primary *FH*-deficient tumors, 12 metastatic *FH*-deficient tumors, 6 HLRCC normal adjacent kidney samples, and 12 normal kidney samples from patients unaffected by HLRCC. Consistent with the functional metabolism results from the HLRCC patient tumor cell lines, mitochondrially encoded genes for the electron transport chain were enriched in normal kidney compared with HLRCC tumors (**Extended Data** Figure 9E). GSEA hallmark analysis showed that while oxidative phosphorylation and fatty acid metabolism were enriched in normal kidneys, glycolysis, EMT, and hypoxia were enriched in the *FH*-deficient tumors (**Fig. 5, G to J, and Extended Data** Figure 10**, A to C**). Consistent with our previous results, highly metastatic and glycolytic *FH*-deficient tumors had an increased EMT signature compared to normal adjacent renal tissue (**Extended Data** Figure 11**, A to C**). Inhibition of oxidative phosphorylation and increased glycolysis, regardless of the mechanism of inhibition, thus promotes EMT and invasion to enable metastasis.

## Discussion

Our findings show that inhibition of oxidative phosphorylation drives a critical transition in the process of metastasis in RCC cancer cells, and that NDUFA4L2 is a modifiable factor that can trigger this shift. By utilizing different implanted models, we observed that NDUFA4L2 inhibits the electron transport chain and OxPhos. OxPhos inhibition was important for triggering a shift in primary tumor cell EMT features and cellular escape, but notably did not drive later steps required for establishing metastatic growth. We further corroborate that inhibition of OxPhos through naturally occurring metabolic mutations, such as loss of *FH*, also linked diminished OxPhos with increased EMT and a proclivity for metastatic spreading. These results, coupled with chemical inhibitor studies, suggest a fundamental paradigm in which the inhibition of OxPhos, regardless of the mechanism of inhibition, facilitates an early step in the transition to metastasis, via metabolically driven changes in chromatin accessibility of EMT genes.

The metastatic process involves metabolic plasticity, with different requirements at discrete phases of metastasis. OxPhos has been proposed to be important to support metastasis in ccRCC patient tumors^10^. Our results add to those findings by highlighting a dynamic set of transitions that take place to facilitate the early steps of the metastatic process. It is likely that cancer cells undergo further shifts in metabolic states in the transition to invasive growth at the metastatic site^2^. Consistent with our findings, it is possible that cells that undergo epithelialization via MET would likely have higher rates of oxidative phosphorylation. Additional lineage tracing studies are needed to determine how metabolism changes as cancer cell progress through the metastatic cascade. Other studies have also suggested high levels of glycolysis may contribute to EMT and metastasis. Hexokinase 2 (HK2), an enzyme that coverts glucose to glucose-6-phosphate, has been reported to increase the expression of SNAIL, a C2H2-type zinc finger transcription factor that represses the epithelial adhesion protein E-Cadherin to promote EMT in breast cancer^26^. This finding is consistent with our data that shows HK2 is increased when cancer cells are in a mesenchymal state and NDUFA4L2 is expressed. The interplay between cell state change and metabolic transitions is likely complex and dynamic, with gene expression changes that alter metabolic signaling providing fundamental cues to enable these transitions to occur.

Although expression of NDUFA4L2 is highest in ccRCC, it is likely that NDUFA4L2 is relevant in other cancers under conditions such as hypoxia. Indeed, NDUFA4L2 has been reported to be an unfavorable clinical prognostic marker in glioblastoma^27^, colorectal cancer^28,29^, and lung cancer^30^. It is also possible that NDUFA4L2 may be widely related to stemness and migration outside of cancer, as NDUFA4L2 was recently found to drive increased stemness and cell viability in human induced pluripotent stem cells (hiPSCs) in response to hypoxia^13^. More studies are needed to identify other contexts in which NDUFA4L2 is relevant and the mechanism in which NDUFA4L2 is expressed. Ultimately, these findings suggest that EMT is facilitated by inhibition of oxidative phosphorylation and that NDUFA4L2 provides is a critical trigger to switch cell states, and reflects a cancer cell specific metabolic target to prevent primary tumor escape and metastasis.

## Materials and Methods

### Patient, UOK365 cell line, Sanger-Sequencing

A 70-year-old male with metastatic HLRCC was evaluated and treated at the Clinical Center of the National Institute of Health on a Urologic Oncology Branch, National Cancer Institute protocol (NCI-97-C-0147) approved by the NCI’s Institutional Review Board and written informed consent was given for participation in this study. Briefly, a fragment of the patient’s tumor was grown as a xenograft in immunodeficient mice, and the resulting tumor was dissociated and contaminating mouse cells removed using magnetic beads (Miltenyi). The resulting tumor cell culture was grown through 10 passages in culture before being designated as the continuous tumor cell line UOK365. FH^-/-^ UOK365 cells were analyzed using both next generation and Sanger mtDNA sequencing with established methods and found to harbor a homoplasmic 19 base pair deletion in the mitochondrial encoded *MT-ND5* gene, resulting in a frameshift mutation (p.Phe478fs) that truncates the *MT-ND5* protein product by 124 residues, or approx. 21% of the total protein length. The *MT-ND5* mutation was also present in DNA obtained from the patient’s tumor at a heteroplasmy level of 73.4%, but was not present in the germline blood DNA.

### Cell Lines

RENCA cells (ATCC CRL-2947) were grown in RPMI supplemented with 10% FBS, 1% penicillin/streptomycin, 1% l-glutamine, 1% HEPES, 1% sodium pyruvate, and 1% MEM amino acid. LVRCC67^16^ cells were grown in Dulbecco’s modified Eagle’s medium and Ham’s F12 mixed 1:1 (Gibco; 11320-033) supplemented with 10% FBS and 1% penicillin–streptomycin, 5 μg ml^−1^ insulin (Sigma, I6634), 1.25 ng ml^−1^ prostaglandin E1 (PGE1; Sigma; P7527), 34 pg ml^−1^ triiodothyronine (T3; Sigma; T5516), 5 μg ml^−1^ apo-transferrin bovine (Sigma; T1428), 1.73 ng ml^−1^ sodium selenite (Sigma; S5261), 18 ng ml^−1^ hydrocortisone (Sigma; H0396), 25 ng ml^−1^epidermal growth factor (Invitrogen; 13247051). All human HLRCC cell lines were grown in DMEM containing high glucose (4.5g/L), supplemented with 10% FBS, 1% penicillin/streptomycin, 1% l-glutamine, and 1% MEM amino acids. All FBS was heat-inactivated before use. Cell lines were subjected to STR and mycoplasma testing to verify purity and identity.

### Mice

BALB/cJ (strain #:000651) mice were purchased from The Jackson Laboratory. Our study exclusively used female mice. It is unknown whether the findings are relevant for male mice. All mouse procedures were carried out under institutional Animal Care and Use Committee (IACUC)-approved protocols from the Vanderbilt University Medical Center (VUMC) and conformed to all relevant regulatory standards. Mice were house in ventilated cages with at most five mice per cage and were provided with ad libitum food and water. Mice were maintained on 12-hr light-dark cycles that coincided with daylight in Nashville, TN. The mouse housing facility was maintained at 20-25°c and 30-70% humidity.

#### Tumor studies

Eight-to twelve-week-old mice were used for injectable tumor models. Mice were euthanized if a humane endpoint was reached (2 cm tumor dimension, ulceration, weight loss >10%). For subcutaneous injection, RENCA cells were trypsinized and washed 3 times in PBS, and 1 × 10^6^ cells were injected in a volume of 100–200 μL PBS into mouse flanks. Subcutaneous tumors grew for 14–21 days. For orthotopic injections, cells were trypsized and washed 3 times in PBS, and 2 × 10^5^ cells were injected in a volume of 100 μL PBS into the subrenal capsule. Orthotopic tumors grew for 18–24 days. For retroorbital injections, cells were trypsized and washed 3 times in PBS, and 1 × 10^5^ cells were injected in a volume of 100 μL PBS into the retro-orbital. Retroorbital tumors grew in the lung for 14–18 days.

### CRISPR editing

The following CRISPR guides were used according to the Zhang Laboratory protocol and single-guide scaffold PX458^32^. For *NDUFA4L2*: forward, CACCGAGATGGCAGGAACTAGTCTA; for reverse, aaacTAGACTAGTTCCTGCCATCTC. Briefly, 1 μg PX458 (Addgene #48138) was incubated with digest-specific reagents for 30 minutes at 37°C. Digested plasmid was gel purified using the QIAquick Gel Extraction Kit (QIAGEN) and eluted prior to phosphorylation and thermocycler-mediated annealing of each pair of oligonucleotides. Ligation reaction was performed per the Zhang protocol using Quick Ligation buffer and Quick Ligase (NEB M2200).

### Lentiviral transduction

NDUFA4L2 and NDI1 3^rd^-generation lentiviral expression vectors were designed with a puromycin resistance cassette and purchased from Vectorbuilder Inc. HEK293T cells were used to package the vectors into lentiviruses. HEK293T cells were cultured until they reached 60% confluence. For each plate, a transfection media cocktail was prepared using a 4:2:1 ratio of Vector of interest, PAX2, and pMD2.G, respectively and Lipofectamine 2000 (ThermoFisher 11668027), according to manufacturer’s instructions. The transfection cocktail was added to the HEK293T cells and incubated for 12 hours at 37°C. Transfection media was then replaced with 10 mL fresh growth media. 48hrs after transfection the lentiviral supernatants were collected and passed through a 0.45 micron filter and were used immediately to infect target cells. Media was replaced on target cells 24hrs after infection. 48-72hrs after transfection, cells were selected using 1.5ug/mL puromycin.

### Protein extraction and western blotting

Cells were harvested and lysed in RIPA buffer supplemented with 1× Halt protease and phosphatase inhibitor cocktail (ThermoFisher 78442) while being kept on ice. Cell lysates were sonicated and pelleted via centrifugation at 14,000*g* for 5 minutes at 4°C, and supernatants were collected in fresh tubes for immediate SDS-PAGE experimentation or preservation at –80°C. Protein concentrations were determined using BCA protein quantification. For SDS-PAGE, 20– 50 μg protein per sample was loaded into and well on a 4%–20% gradient polyacrylamide gel (Bio-Rad), followed by transfer onto a nitrocellulose membrane. The membranes were then subjected to immunoblotting with the primary antibodies E-Cadherin (1:1000, Cell Signaling, 3195) and β-actin (1:1,000, MilliporeSigma, A2066) was used as a loading control.

#### Native Immunoblots

The NativePAGE Novex Bis-Tris gel system (Thermo Fisher Scientific) was used to analyze native membrane protein complexes and native mitochondrial matrix complexes, with the following modifications: only the Light Blue Cathode Buffer was used; 20 μg of membrane protein extracts were loaded/well; the electrophoresis was performed at 150 V for 1 hour and 250 V for 2 hours. For the native immunoblot, PVDF was used as the blotting membrane. The transfer was performed at 25 V for 4 hours at 4 °C. After transfer, the membranes were washed with 8% acetic acid for 20 min to fix the proteins, and then rinsed with water before air-drying. The dried membranes were washed 5-6 times with methanol (to remove residual Coomassie Blue G-250), rinsed with water and then blocked for 2 hours at room temperature in 5% milk, before incubating with the desired antibodies diluted in 2.5% milk overnight at 4 °C.

### 3D Invasion Assay

Growth factor reduced Matrigel (Corning, 10–12 mg/mL stock concentration, catalog no. 354230) and rat tail (Corning, catalog no. 354236) Collagen I were used for organotypic culture experiments. Vertical invasion assays and experiments in three-dimensional (3D) culture were performed and quantified as described previously^33^ using a Matrigel/Collagen I matrix (3–5 mg/mL Matrigel and 1.8–2.1 mg/mL Collagen I). Slides were stained using Phalloidin and Hoechst (**Extended Data Table 1**) and visualized with confocal microscopy using a Zeiss LSM 880.

### Histology

Tumors were harvested and fixed with 10% neutral buffered formalin solution for 48 hours on a shaker at room temperature. Tissues were then washed in PBS and embedded in paraffin for sectioning by the Translational Pathology Shared Research Core at VUMC. All tissues were sectioned at 5 μm. After serial sectioning, H&E was performed on each tissue and IHC was performed as previously described^34^. Briefly, slides were deparaffinized in xylene and rehydrated in serial ethanol dilutions. Antigen retrieval was performed by heating slides for 17 minutes in Tris EDTA buffer, pH9, in a pressure cooker at 110°C. Slides were cooled and blocked with 2.5% horse serum (Vector Laboratories). After blocking, slides were incubated overnight at 4°C with a primary antibody in horse serum **(Extended Data Table 1)** . Slides were then incubated in anti-rabbit HRP secondary (Vector Laboratories) for 1hr at room temperature the following day and subsequently incubated in 1:500 Opal 520 (Akoya FP1487001KT) or Opal 570 (Akoya FP1488001KT) for 10 minutes. For serial staining, slides were stripped using Citric Acid buffer, pH 6.1 in a pressure cooker at 110°c for 2min and then staining was repeated using different antibody and Opal fluorophore. After last Opal staining, slides were mounted using antifade gold mount with DAPI (Invitrogen). Stained images were acquired using an Aperio Versa 200 Automated Slide imaging system (Leica/Aperio) via the Vanderbilt University Medical Center Digital Histology Shared Resource (DHSR) core at 20X magnification. Images were analyzed with Fiji software. Quantification of markers was done by measuring total amount of fluorescence divided by total area of tissue (H&E).

### Extracellular flux assays

#### ccRCC patient tumors

Mechanical dissociation of human tumors using Miltenyi gentleMACS (human setting) and enzymatic digestion, as described above, were performed prior to sequential isolations using Miltenyi bead isolation kits (CD45, #130-052-301). The CD45 negative isolated fraction was plated at 200,000 live cells/well in 4–8 technical replicates on a Cell-Tak–coated plate (Corning, 354240) in Agilent Seahorse RPMI 1640 supplemented with 10 mM glucose, 1 mM sodium pyruvate, and 2 mM glutamine. Live cells were used for RENCA (50,000), LVRCC67 (50,000), and HLRCC human cell lines (7,000) Seahorse flux assays. Cell metabolism was analyzed on a Seahorse XFe 96 bioanalyzer using the Mitostress assay (Agilent Technologies, 103015-100) with 1μM oligomycin, 2 μM FCCP, and 0.5 μM rotenone/antimycin A. Measurements were normalized using total cell counts, measured from flow cytometry taken directly after the seahorse assay. Data were analyzed with Agilent Wave software, version 2.6.

### Flow Cytometry and SCENITH

#### DCFDA, Mitotracker, TMRE, and MitoSox analysis

Cells were stained for flow cytometry analysis as previously described^35^ using antibodies in **Extended Data Table 1**. Cells were washed and stained and data were analyzed using FlowJo (BD Biosciences) with mean fluorescence intensity (MFI) calculated using the geometric mean.

#### Tumor and spleens

Mice were euthanized and tumors were collected as previously described^15^. Single-cell suspensions of splenocytes were prepared by mechanical dissociation followed by ACK lysis. Tumors were chopped, mechanically dissociated on the Miltenyi gentleMACS Octo Dissociator with Heaters (setting implant tumour one) and digested in 435 U ml^−1^ DNase I (Sigma-Aldrich, D5025) and 218 U ml^−1^ collagenase (Sigma-Aldrich, C2674) at 37 °C for 30 min. After enzyme treatment, tumors were passed through a 70-μm filter and ACK-lysed. Cells were resuspended in complete RPMI and counted using trypan blue with the TC20 Automated Cell Counter (Bio-Rad). Tumor cells were stained for flow cytometry analysis a previously described using antibodies in **Extended Data Table 1**^15^. Data were analyzed using FlowJo (BD Biosciences) with mean fluorescence intensity (MFI) calculated using the geometric mean.

#### SCENITH

RENCA cells were seeded in 96 well plates with technical triplicates for each experiment and SCENITH was performed as previously described^17^. Briefly, wells were treated for 30-45 minutes with DMSO, 2-Deoxy-D-Glucose (2DG, final concentration 100mM), Oligomycin (Oligo, final concentration 1μM), or a sequential combination of the drugs at the final concentrations previously mentioned. Puromycin (final concentration 10 μg/ml) was added during the last 15-45 minutes of the treatment with metabolic inhibitors. After puromycin treatment, cells were washed in cold PBS and stained with a combination of Fc receptors blockade and fluorescent cell viability marker, then primary conjugated antibodies against surface markers listed in **Extended Data Table 1**, during 25 minutes at 4°C in PBS with 5% FBS (FACS buffer). After washing, cells were fixed and permeabilized using FOXP3 fixation and permeabilization buffer (Thermofisher eBioscience™) following manufacturer instructions.

Intracellular staining of puro was performed by incubating cells during 1 hour at 4°C diluted in permeabilization buffer. Data were analyzed using FlowJo (BD Biosciences) with mean fluorescence intensity (MFI) calculated using the geometric mean. Glycolytic capacity is defined as the maximum capacity to sustain protein synthesis levels when mitochondrial OxPhos is inhibited. The percentage of glucose dependence quantifies how much the translation levels are dependent on glucose oxidation and is calculated as the difference between puro levels in 2-Deoxy-D-Glucose treated cells compared to control, divided by the difference in puro upon complete inhibition of ATP synthesis (2DG, first and then Oligomycin A, combined) compared to control cells. Percent of mitochondrial dependence quantifies how much translation is dependent on oxidative phosphorylation and is defined as the difference in puro levels in Oligomycin A treated cells compared to control relative to the decrease in puro levels upon full inhibition of ATP synthesis inhibition (combination of 2DG and Oligomycin) also compared to control cells.

### Colony Formation

To assess the impact of the experimental conditions on cell growth, 50 cells were seeded into 6-well plates, respectively, and allowed to adhere for 24 hours. On the following day, growth medium was supplemented with vehicle or various doses of 2-Deoxyglucose (2DG), and the effect on growth was monitored after two weeks by staining with a 1% crystal violet solution prepared in 20% methanol and 80% dH_2_O. After imaging, the crystal violet stain was washed with water and total area of crystal violet was quantified using Image J.

### Electron Microscopy

#### Cell Lines

All electron microscopy reagents were purchased from Electron Microscopy Sciences. Cell cultures were grown to approximately 80% confluence and fixed with 2.5% glutaraldehyde in 0.1 M Na cacodylate buffer for 1 hour at room temperature followed by 24 hours at 4°C. After fixation the cells were mechanically lifted from the tissue culture plates and pelleted, then sequentially postfixed with 1% tannic acid, 1% Osmium Tetroxide, and stained en bloc with 1% uranyl acetate for 1 hour each. Samples were dehydrated in a graded ethanol series 25%,50%, 70%, 80%, 85%, 90%, and three-100% steps for 15 minutes each. Following dehydration the samples were infiltrated with Quetol 651–based Spurr’s resin composed of Quetol 651, ERL 4221, NSA, DER 736, and BDMA (Electron Microscopy Sciences) at 50%, 75%, and three-100% resin exchanges using propylene oxide as the transition solvent. Samples were polymerized in Eppendorf tubes at 60°C for 48 hours. Ultrathin sections were prepared on a Leica UC7 ultramicrotome at a nominal thickness of 70 nm and collected onto 300-mesh nickel grids. Sections were stained with 2% uranyl acetate and lead citrate. Cell samples were imaged using a Tecnai T12 operating at 100 kV equipped with an AMT NanoSprint CMOS camera using AMT imaging software.

#### HLRCC Patient Samples

Tissue samples were fixed with 2% glutaraldehyde and 4% formaldehyde in 0.1M Na Cacodylate buffer for an hour at room temperature for Transmission electron microscopy (TEM) processing. Each tissue sample was cut into five smaller pieces and transferred to a glass vial containing 0.1M Na Cacodylate buffer for EM processing. Tissue samples were washed with 0.1M Na Cacodylate buffer two times for 10 min. After the second buffer wash, the tissue samples were post fixed with 1% Osmium Tetroxide for an hour in the dark. Tissue samples were washed with 0.1M Na Cacodylate buffer twice for 10 min and washed once with 0.1N Na Acetate buffer and then stained with 0.5% Uranyl Acetate for an hour in the dark. After en block staining, tissue samples were washed with 0.1N Na Acetate buffer twice for 10 min. After the last buffer wash, tissue samples were subjected to gradual dehydration in the order of 35%, 50%, 70% and 95% Ethanol twice each step for 10 min and three times with 100% Ethanol for 10 min. After the last step of 100% Ethanol rinse, tissue samples were further dehydrated in Propylene oxide (PO) for 10 min for three times. After the last step in PO, tissue samples were infiltrated in 50:50 epoxy resin and PO overnight at room temperature.

The next day, after overnight infiltration, each tissue sample was removed from 50:50 epoxy resin and PO, blotted, and embedded in a plastic mold containing 100% pure epoxy resin and transferred to a 55 °C oven for 48 hours. Epoxy resin (PolyScience Resin) ingredients consisted of a mixture of Poly/Bed 812 embedding Media, Dodecenylsuccinic Anhydride (DDSA), Nadic Methyl Anhydride (NSA) and DMP-30 to solidify the resin. After 48 hours, tissue samples were taken out of the oven, and the best tissue embedded in resin mold was selected, then ultra-thin sectioned with a UC6 Leica Microtome. The ultra-thin sections were picked up on a 150 Copper mesh grid and were examined under a Hitachi H7600 Transmission electron microscope. The grids were post stained with 1:1 0.5% Uranyl Acetate in ddH2O and 70% Ethanol for 2 min and then rinsed with ddH2O for four times. Then cells stained with 1:1 Lead Citrate and H2O for 2 min and rinsed with ddH2O four times. The grids were carbon coated with a TedPella/Cressington Evaporator and imaged in a Hitachi H7600 transmission electron microscope at 80KeV.

### Targeted Metabolomics

RENCA cells (EV, NDUFA4L2 KO, and NDUFA4L2 Rescue) were grown to 80% confluent. Once at 80% confluency, media was removed, and 100 mm plates were washed twice with room temperature PBS. After washing, 3 mL of -80°C 80:20 MeOH:H_2_O was added to the plates. Culture plates were then placed in -80°C freezer to extract for 15 minutes. Plates were then placed on dry ice and cells were scraped to harvest and added to a 15 mL conical tube containing 100 nmol ^13^C-1-Lactate as an internal standard. The plates were then washed with 3 mL ice cold 80:20 MeOH:H_2_O and the samples were added to the 15 mL conical tubes containing the internal standard. Samples were pelleted by centrifugation at 2,800 x *g* for 10 minutes at 4°C. Supernatant containing extracted metabolites and internal standard was transferred to new 15 mL conical tubes and dried under nitrogen. Precipitated protein pellets were resolubilized and protein concentration was measured via BCA assay (Thermo Fisher Scientific, Waltham, MA). Samples were resuspended in 80 µL 3:2 mobile buffer A: mobile buffer B (see below). 18 µL of the sample was then chromatographed with a Shimadzu LC system equipped with a 100 x 2.1mm, 3.5μm particle diameter XBridge Amide column (Waters, Milford, MA). Mobile phase A: 20 mM NH_4_OAc, 20 mM NH_4_OH, 5% acetonitrile in H_2_O, pH 9.45; mobile phase B: 100% acetonitrile. With a flow rate of 0.45 mL/min the following gradient was used: 2.0 min, 95% B; 3.0 min, 85% B; 5.0 min, 85% B; 6.0 min, 80% B; 8.0 min, 80% B; 9.0 min, 75% B; 10 min, 75% B; 11 min, 70% B; 12 min, 70% B; 13 min, 50% B; 15 min, 50% B; 16 min 0% B; 17.5 min, 0% B; 18 min, 95% B. The column was equilibrated for 3 minutes at 95% B between each sample. Scheduled MRM was conducted in negative mode with a detection window of 120 seconds using an AB SCIEX 6500 QTRAP with the analyte parameters below. All analytes (**Extended Data Table 2**) were quantified via LC-MS/MS using the ^13^C-1-Lactate internal standard and normalized to the protein in each respective sample’s cell pellet. Outliers were removed using interquartile range.

### PTM analysis

#### Chromatin Isolation

Renca cell pellets (EV, NDUF4AL2 KO, NDUF4AL2 Rescue) from 10 cm culture dishes were gently resuspended by pipetting in 300 µL of hypotonic lysis buffer (10 mM HEPES, 1.5 mM MgCl_2_, 10 mM KCl, 0.5% IGEPAL, 5 mM NaB, pH 7.9) containing Halt™ Protease and Phosphatase Inhibitor Cocktail (1:100 v/v, Thermo Fisher Scientific, Waltham, MA) and incubated on ice for 30 minutes. The nuclear fraction was then isolated via centrifugation at 1,000 x *g* for 10 minutes at 4°C. The supernatant, containing the non-nuclear fraction, was removed and protein was measured via BCA assay (Thermo Fisher Scientific, Waltham, MA). The nuclear pellets were then resuspended in a high salt buffer (20 mM HEPES, 1.5 mM MgCl_2_, 420 mM KCl, 0.2 mM EDTA, 25% glycerol, 5 mM NaB, pH 7.9) containing protease and phosphatase inhibitors (1:100 v/v) and rotated end-over-end overnight at 4°C. Chromatin was then isolated via centrifugation at 1,000 x *g* for 20 minutes at 4°C and high salt buffer was removed. Chromatin was resuspended in 200 µL ddH_2_O and sonicated into solution. Concentration of resolubilized chromatin was measured via BCA assay (Thermo Fisher Scientific, Waltham, MA).

#### Quantification and Analysis of Arginine (R) and Lysine (K) Modifications (QuARKMod)

Lys and Arg PTMs (**Extended Data Table 3**) were quantified in EV, NDUF4AL2 KO, NDUF4AL2 Rescue RENCA cell chromatin (see above) and whole cell lysates as described ^36,37^. Briefly, protein from whole cell or nuclear fraction samples were precipitated in ice-cold acetone at -20°C for 1 hour. Next, precipitated protein was pelleted via centrifugation at 16,000 x g for 10 minutes at 4°C. Supernatant was removed and samples were allowed to air dry. Once dry, protein pellets were resuspended in 65 µL of freshly made 50 mM NH4HCO3 (pH 8.0); followed by 10 µL of an internal standard mix (see table below) and 10 µg of sequencing grade trypsin (10 µL, Promega, Madison, WI). Proteins were then digested to peptides via tryptic digest for 16 hours at 37°C. After digestion, trypsin was denatured via boiling at 95°C for 10 minutes. Once samples were had cooled to room temperature, samples were briefly spun on a tabletop centrifuge and 15 µg of aminopeptidase M (10 µL, MilliporeSigma, Burlington, MA) was added to each sample. Peptides were then digested to single amino acids with aminopeptidase M for 16 hours at 37°C. Following digestion, aminopeptidase M was denatured via boiling at 95°C for 10 minutes and samples were cooled to room temperature. 15 µL of a 1:1 solution of heptafluorobutyric acid (HFBA, MilleporeSigma, Burlington, MA) was added to each sample, and samples were spun at 16,000 x g for 10 minutes to remove any insoluble debris. TO measure methylated Arg (ADMA, SDMA, MMA), a standard curve was generated using an ADMA natural isotope standard against the Arg internal standard. 12 µL of supernatant was then chromatographed with a Shimadzu LC system equipped with a 150 x 2.1mm, 3.5 μm particle Eclipse XDB-C8 column (Agilent, Santa Clara, CA) with mobile phase A: 10 mM HFBA in H2O and mobile phase B: 10 mM HFBA in acetonitrile. With a flow rate of 0.325 mL/min the following gradient was used: 0.5 min, 2.5% B; 5.5 min, 50% B; 6.0 min, 80% B; 9 min, 80% B; 9.5 min, 2.5% B. The column was equilibrated for 3 minutes at 2.5% B between samples. MRM was conducted in positive mode using an AB SCIEX 6500 QTRAP. Samples were corrected with enzyme only controls and normalized to Leu abundance in each respective sample. Outliers were removed using interquartile range.

### Single cell RNAseq data processing from GSE178481

Filtered counts for human single cell RNA sequencing data of clear cell renal cell carcinoma were accessed from the NCBI Gene Expression Omnibus database under accession code GSE178481^14^. Counts were loaded into Seurat v5.1^38^ and merged into a single object using sample and cell annotations provided by the original authors. Data were normalized using the *NormalizeData* function and the top 3000 variable features were selected for dimensional reduction using the *FindVariableFeatures* function with default parameters. For dimensional reduction, batch effects were modeled using Harmony^39^ with a *batch_key* corresponding to individual patient identifiers and uniform manifold approximation and projection (UMAP) was calculated using the *RunUMAP* function with *min.dist=0.5* and *spread=1.* For downstream analyses or plotting requiring Scanpy^40^ objects, scDIOR ^41^ was used for object conversion.

#### Single cell pathway scoring

The scMetabolism^31^ package was used to estimate pathway activity for KEGG metabolic pathways using the AUCell^42^ method. The Wilcoxon rank sum test was used to detect differential activity of metabolic pathways with the *FindMarkers* function in Seurat v5.1 and the fast implementation by Presto^43^. UCell^44^ v.2.6 was utilized to score the hypoxia hallmark gene set from the Molecular Signatures Database^45^.

### Single cell RNAseq data processing for fig. S1

#### Data Acquisition and Preprocessing

The raw single-cell RNA sequencing (scRNA-seq) data were obtained from PMID: 33711272, and accessed via the Single Cell Portal (https://singlecell.broadinstitute.org/single_cell/study/SCP1288/tumor-and-immune-reprogramming-during-immunotherapy-in-advanced-renal-cell-carcinoma#study-summary)^11^. Initial filtering was applied to retain high-quality cells by removing cells with high mitochondrial content or low gene counts, which could indicate damaged cells, doublets, or multiplets. Cells with more than 3,000 detected genes were excluded to remove damaged cells or potential doublets and multiplets. Cells with mitochondrial gene expression contributing more than 5% of total counts were removed to exclude cells likely to be under stress or undergoing apoptosis.

#### Normalization and Transformation

To account for differences in sequencing depth across cells, the filtered data were normalized. Total counts for each cell were scaled to 10,000 counts, followed by a log transformation to stabilize variance and approximate a normal distribution. Counts were normalized to 10,000 counts per cell to control for differences in sequencing depth. A natural log transformation (log1p) was applied, where each count was transformed as log(count + 1), stabilizing variance and enabling downstream statistical analysis. Additional filtering criteria were applied to focus on biologically meaningful genes and cells. Cells with fewer than 200 detected genes and genes expressed in fewer than 3 cells were removed.

#### Visualization of Gene Expression Patterns

To visualize gene expression patterns across cell lineages, a UMAP projection was generated using the dataset prior to subsetting for differential expression analysis. Cells were colored by lineage to observe clustering patterns associated with different cellular lineages, including the Putative Tumor subset that was later used in downstream analyses. To specifically visualize SNAI2 expression in the UMAP, point sizes were adjusted based on SNAI2 expression levels. Cells with detectable SNAI2 expression were represented with larger points to enhance visibility and highlight cells expressing SNAI2.

### RNA isolation, qPCR, and RNA-seq analyses

#### Cell lines

Total RNAs were isolated and purified using the RNeasy Mini Kit (QIAGEN, 74106). Isolated RNA was converted to cDNA using iScript Reverse Transcription Supermix (Bio-Rad). mRNA expression was measured with a real-time PCR detection system (Applied Biosystems) in 96-well optical plates using SsoAdvanced Universal SYBR Green Supermix (Bio-Rad). β-Actin was used as a control. All primers utilized are listed in **Extended Data Table 4**.

#### RENCA scRNA-seq sample processing

10-week-old female BALB/c mice were injected subcutaneously with 1 × 10^6^ WT RENCA cells. Tumors were harvested 16 days after injection and then enzymatically digested as described above. CD45^+^ and CD45^−^ live cells were flow sorted and counted using trypan blue and a Bio-Rad TC20 counter. Three tumors of approximately the same weight were pooled together. And CD45^+^ hashing antibodies were applied individually. Flow-sorted cells were incubated with FcX block for 10 minutes at 4°C and then washed with cell staining buffer. Samples were then pooled and resuspended at 500,000 cells/mL in PBS plus 0.4% BSA with a target of 10,000 live cells loaded onto the Chromium Controller (10x Genomics) and processed according to the manufacturer’s instructions. Sequencing was performed on the Illumina NovaSeq 6000, targeting 50,000 reads per cell for the 5′ assay. The raw data (FASTQ files) were demultiplexed and processed using CellRanger (version 6.1.2) to get the gene expression matrix with the reference genome mm10 by VANTAGE. 10X genomics software was used to evaluate the gene expression of EMT markers in CD45-cells. Raw data was published in Wolf et al.^15^ and was deposited in the Gene Expression Omnibus (GEO) database (GSE239889).

#### HLRCC patient tumors RNA-seq analysis

Total RNA was extracted from cell lines or cryo-pulverized tumor/normal tissues using Trizol Reagent (Invitrogen) following the manufacturer’s protocol and stored at −80 °C. RNA concentrations and purities were measured using a NanoDrop 2000 UV-Vis Spectrophotometer (Thermo Fisher Scientific) and 5μg of total RNA was sent to ACGT, Inc. (Germantown, MD, USA) for RNAseq analysis. All samples were processed with the TruSeq mRNA kit using oligo-dT beads to capture polyA tails and data was delivered as a Multiplex Fast-Q file with adaptors trimmed. 43-60 million read pairs per sample were obtained. The sequencing quality of the reads was assessed using FastQC (version 0.11.5), Preseq (version 2.0.3)^46^, Picard tools (version 1.119), and RSeQC (version 2.6.4). Reads were trimmed using Trimmomatic (version 0.33)^47^ to remove sequencing adapters, prior to mapping to the human reference genome, hg38, using STAR (version 2.5.2b) in two-pass mode^48^. Across the samples, the median percentage of mapped read pairs was 76.8%. Expression levels were quantified using RSEM version 1.3.0^49^ with GENCODE annotation version 21^50^. The RSEM counts were then normalized using the variance stabilization transformation^51^ from the DESeq2 R package (version 1.24.0)^52^. These normalized values were used for visualizing mRNA expression levels and pathway enrichment.

### ATAC-sequencing via Novogene

ATAC-seq was performed as previously reported^53–55^ by Novogene Co, Ltd. Briefly, nuclei were extracted from samples, and the nuclei pellet was resuspended in the Tn5 transposase reaction mix. The transposition reaction was incubated at 37°C for 30 min. Equimolar Adapter1 and Adatper 2 were added after transposition, PCR was then performed to amplify the library. After the PCR reaction, libraries were purified with the AMPure beads and library quality was assessed with Qubit.

The clustering of the index-coded samples was performed on a cBot Cluster Generation System using TruSeq PE Cluster Kit v3-cBot-HS (Illumina) according to the manufactuer’s instructions. After cluster generation, the library preparations were sequenced on an Illumina Hiseq platform and 150 bp paired-end reads were generated.

Nextera adaptor sequences were firstly trimmed from the reads using skewer (0.2.2). These reads were aligned to a reference genome using BWA, with standard parameters. These reads were then filtered for high quality (MAPQ≥13), non mitochondrial chromosome, and properly paired reads (longer than 18 nt).

All peak calling was performed with macs2 using ‘macs2 callpeak --nomodel --keepdup all --call-summits’. For simulations of peaks called per input read, aligned and de-duplicated BAM files were used without any additional filtering. ATAC-seq graphs were produced using https://igv.org/app/ with the mouse genome (mm10) used as reference.

### Cell Line Bulk RNA-sequencing and Analysis via Novogene

#### Library Construction, Quality Control, and Sequencing

RNA was isolated from RENCA using Qiagen RNeasy Plus Kit per manufacturer’s instructions. RNA was sent to Novogene to perform library prep, quality control, and sequencing. Briefly, messenger RNA was purified from total RNA using poly-T oligo-attached magnetic beads. After fragmentation, the first strand cDNA was synthesized using random hexamer primers, followed by the second strand cDNA synthesis using either dUTP for directional library or dTTP for non-directional library^56^. For the non-directional library, it was ready after end repair, A-tailing, adapter ligation, size selection, amplification, and purification.

For the directional library, it was ready after end repair, A-tailing, adapter ligation, size selection, USER enzyme digestion, amplification, and purification. The library was checked with Qubit and real-time PCR for quantification and bioanalyzer for size distribution detection. Quantified libraries will be pooled and sequenced on Illumina platforms, according to effective library concentration and data amount.

#### Bioinformatics Analysis

Raw data (raw reads) in fastq format was firstly processed through in-house perl scripts. In this step, clean data (clean reads) was obtained by removing reads containing adapter, reads containing ploy-N and low quality reads from raw data. At the same time, Q20, Q30 and GC content were calculated. All the downstream analyses were based on the clean data with high quality. Reference genome and gene model annotation files were downloaded from genome website directly. Index of the reference genome was built using Hisat2 v2.0.5 and paired-end clean reads were aligned to the reference genome using Hisat2 v2.0.5. We selected Hisat2^57^ as the mapping tool for that Hisat2 can generate a database of splice junctions based on the gene model annotation file and generate a better mapping result than other non-splice mapping tools. The mapped reads of each sample were assembled by StringTie^58^ (v1.3.3b) in a reference-based approach. StringTie uses a novel network flow algorithm as well as an optional de novo assembly step to assemble and quantitate full length transcripts representing multiple splice variants for each gene locus.

FeatureCounts^59^ v1.5.0-p3 was used to count the reads numbers mapped to each gene. Then FPKM of each gene was calculated based on the length of the gene and reads count mapped to this gene. FPKM, expected number of Fragments Per Kilobase of transcript sequence per Millions base pairs sequenced, considers the effect of sequencing depth and gene length for the reads count at the same time, and is currently the most commonly used method for estimating gene expression levels. Differential expression^60^ analysis of two conditions/groups (two biological replicates per condition) was performed using the DESeq2R package (1.20.0). DESeq2 provide statistical routines for determining differential expression in digital gene expression data using a model based on the negative binomial distribution. The resulting P-values were adjusted using the Benjamini and Hochberg’s approach for controlling the false discovery rate. Genes with an adjusted P-value <=0.05 found by DESeq2 were assigned as differentially expressed. (For edgeR (Robinson, 2010 #1411) without biological replicates) Prior to differential gene expression analysis, for each sequenced library, the read counts were adjusted by edgeR program package through one scaling normalized factor. Differential expression analysis of two conditions was performed using the edgeR R package (3.22.5). The P values were adjusted using the Benjamini & Hochberg method. Corrected P-value of 0.05 and absolute foldchange of 2 were set as the threshold for significantly differential expression.

Gene Ontology (GO) enrichment analysis of differentially expressed genes was implemented by the clusterProfiler R package, in which gene length bias was corrected. GO terms with corrected Pvalue less than 0.05 were considered significantly enriched by differential expressed genes. KEGG is a database resource for understanding high-level functions and utilities of the biological system, such as the cell, the organism and the ecosystem, from molecular-level information, especially large-scale molecular datasets generated by genome sequencing and other high-through put experimental technologies (http://www.genome.jp/kegg/). We used clusterProfiler R package to test the statistical enrichment of differential expression genes in KEGG pathways. Gene Set Enrichment Analysis (GSEA) is a computational approach to determine if a pre-defined Gene Set can show a significant consistent difference between two biological states. The genes were ranked according to the degree of differential expression in the two samples, and then the predefined Gene Set were tested to see if they were enriched at the top or bottom of the list. Gene set enrichment analysis can include subtle expression changes. We use the local version of the GSEA analysis tool http://www.broadinstitute.org/gsea/index.jsp, GO, KEGG data set were used for GSEA independently.

### Statistics

GraphPad Prism 10 (GraphPad Software) was used to create graphs and conduct statistical analyses. Data are expressed as the mean ± SD. Analyses of the differences between 2 test groups were performed using a nonparametric, unpaired, 2-way *t* test. For analysis of 3 or more groups, a 1-way ANOVA was performed with Dunnett’s or Bonferroni’s test. *P* values were considered statistically significant if *P* was less than 0.05.

### Study approval

Fresh, histology-confirmed ccRCC tumors were surgically removed from 7 patients included in this study. All studies were conducted in accordance with Declaration of Helsinki principles under a protocol approved by the VUMC IRB (protocol 151549). Informed consent was received from all patients prior to inclusion in the study by the Cooperative Human Tissue Network at VUMC. HLRCC patient samples were collected using National Cancer Institute protocol (NCI-97-C-0147) approved by the NCI’s Institutional Review Board and written informed consent was given for participation in this study. All mouse procedures were implemented under VUMC IACUC-approved protocols and conformed to all relevant regulatory standards.

## Acknowledgments

We thank members of the laboratories of J.C.R. and W.K.R. for input; We also thank A. Hakimi for his generous gift of LVRCC67 cells. All schematics were created using Biorender.com. We acknowledge the Translational Pathology Shared Resource supported by NCI/NIH Cancer Center Support Grant P30CA068485. Flow Cytometry experiments were performed in the VUMC Flow Cytometry Shared Resource. Some electron microscopy was performed at the Vanderbilt Cell Imaging Shared Resource. The VUMC Flow Cytometry Shared Resource and the Vanderbilt Cell Imaging Shared Resource are supported by the Vanderbilt Ingram Cancer Center (P30 CA68485) and the Vanderbilt Digestive Disease Research Center (DK058404). We also thank the VUMC Digital Histology Shared Resource for whole slide imaging. The current affiliation of M.L. is Institute for Biomedicine, Eurac Research, Bolzano, Italy. This study was funded by National Institutes of Health grant K00 CA253718 (ENA), National Institutes of Health grant F31 CA261049 (MMW), F31 DK139660, National Institutes of Health grant T32 CA009582 (EQJ), National Institutes of Health grant R01 CA217987 (JCR), National Institutes of Health grant R01 DK105550 (JCR), Department of Defense W81XWH-22-1-0419 (JCR), Department of Defense KC210152 (WKR and JCR), Department of Defense W81XWH2110786 (FMM), National Institutes of Health grant K12 CA090625 (KEB and WKR), The Vanderbilt-Incyte Alliance (JCR and WKR), The Mark Foundation for Cancer Research (JCR and KI), National Cancer Institute grant ZIA BC011028 (WML), and National Cancer Institute grant ZIC BC011932 (DRC and WML).

## Author contributions

ENA, WKR, and JCR conceptualized the study. ENA, EQJ, MMW, LV, MAC, ELFG, EK, KKS, AS, ZH, AB, DC, CJR, ML, NM, YS and CJR contributed experimentally. WKR and JCR obtained funding. KEB, FMM, WML, KI, WKR, JCR supervised the study. ENA wrote the original draft. ENA, WKR, and JCR reviewed and edited the manuscript.

## Data and materials availability

All data including the sequencing data included in this study are available in the main text, supplementary materials, or publicly available. The bulk RNA-sequencing, ATAC-sequnecing, and HLRCC RNA-seq will be deposited in the Gene Expression Omnibus (GEO) da tabase (pending). P ublicly available databases used for this study include GSE239889 (RENCA scRNA-seq), G SE178481 (Fig. 1), a nd dbGaP: phs002065.v1.p1(Extended Data Figure 1). NDUFA4L2 expression from the TCGA database were accesssed using Kmplot.com ^61^. UOK365 and LVRCC67 cells were recived with MTA from NCI and MSKCC, respectively.

Supplementary information is available for this paper.

Correspondence and requests for materials should be addressed to Jeffrey Rathmell at.

## Supplementary Materials

Extended Data Figures 1 to 11

Extended Data Tables 1-4

References *1*–*61* (1-30 are in main text)

