## Supplemental Figures and Tables for "Impaired oxidative phosphorylation drives primary tumor escape and metastasis"

**Running Title:** Inhibition of OxPhos drives EMT

**The PDF file includes:**

Extended Data Figures 1 to 11
Extended Data Tables 1 to 4

### Extended Data Figures

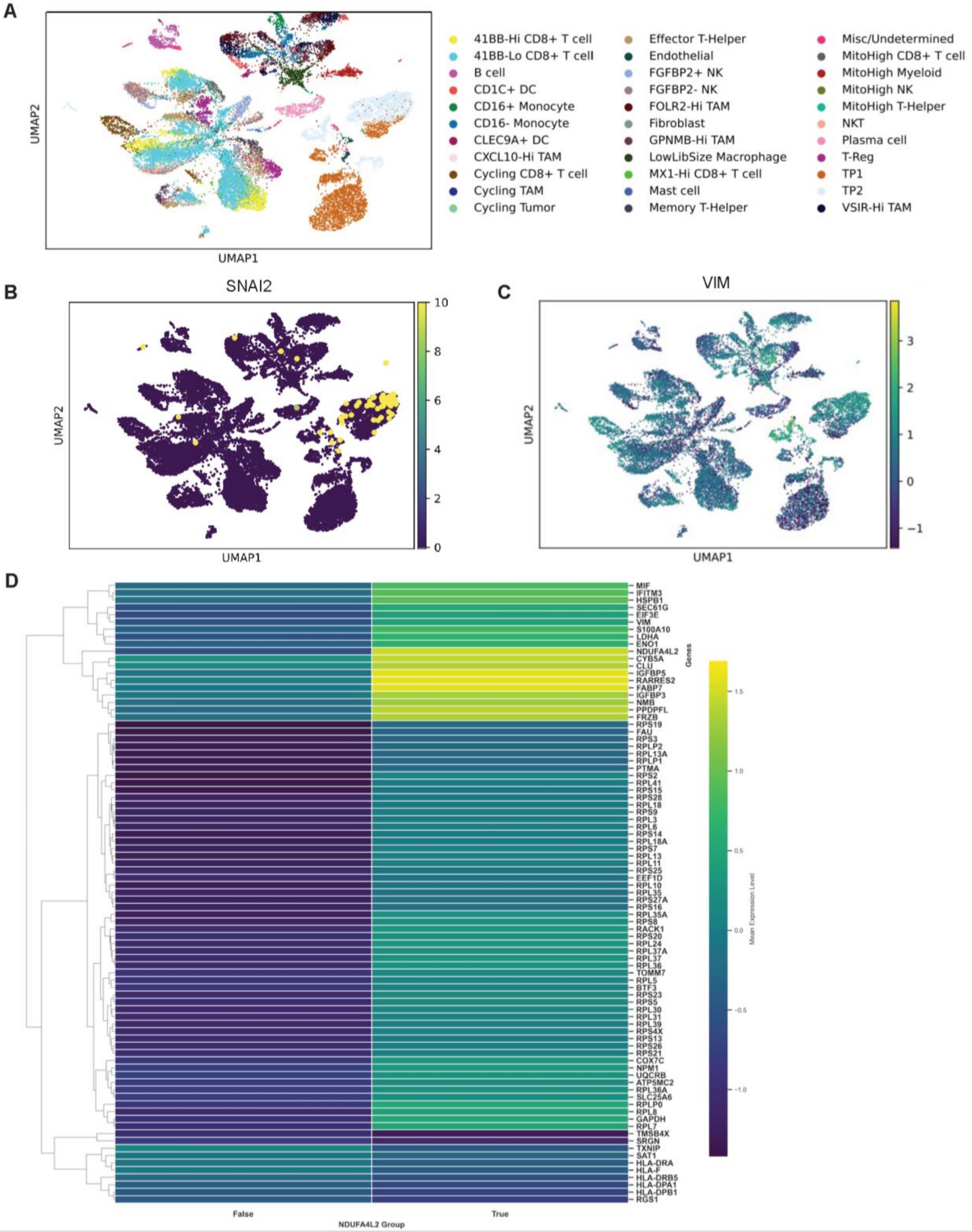

**Extended Data Figure 1. EMT is correlated with increased glycolysis in RCC. (A)** Single-cell RNA sequencing expression clustering from Bi et al[11]. Expression of mesenchymal markers *SNAI2* **(B)** and *VIM* **(C)** were found in the tumor population TP2, as defined by the original authors. The TP2 tumor cell population is associated with increased glycolysis and VHL loss-of-function mutations. **(D)** Differential gene expression between *NDUFA4L2* negative (false) and positive (true) tumor cells using a Wilcoxon rank-sum test from (A).

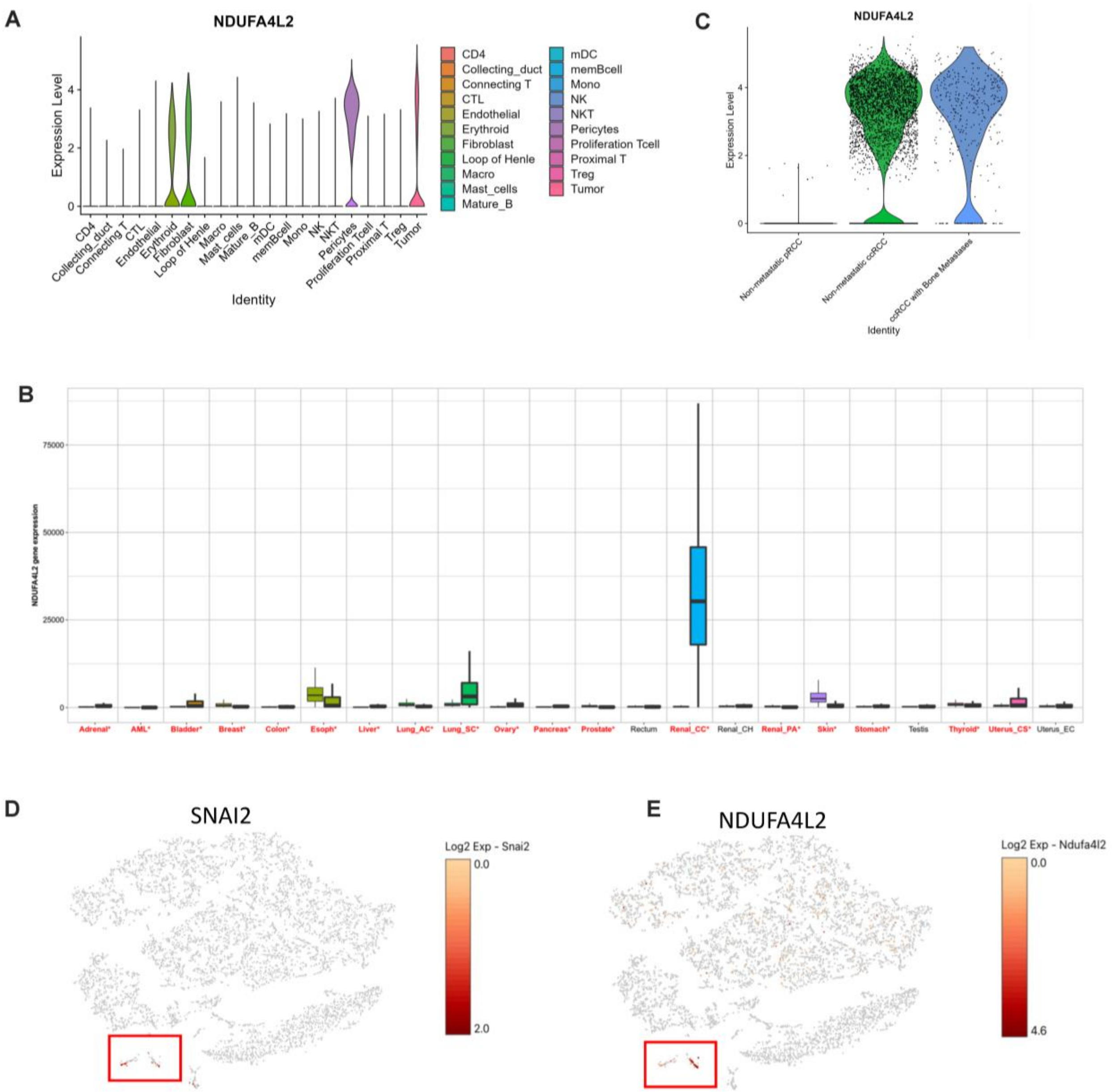


**Extended Data Figure 2. NDUFA4L2 is correlated with EMT. (A)** In scRNA-seq from GEO accession #GSE178481[14], expression of *NDUFA4L2* was largely found in the tumor cells, pericytes, fibroblasts, and erythroid cells, but not immune cells. Cell annotations from original authors. **(B)** Bioinformatic analysis of *NDUFA4L2* mRNA expression in TCGA pan-cancer database, identified *NDUFA4L2* as elevated in ccRCC tumor tissue, but not normal kidney. **(C)** Also in the scRNA-seq analysis from (A), *NDUFA4L2* expression was high in tumors with non-metastatic ccRCC and metastatic ccRCC but not papillary RCC tumors. Single-cell RNA sequencing was done on CD45 negative cells isolated from syngenic subcutaneous RENCA RCC tumors and analyzed using 10X genomics software. **(D)** *Snai2* and **(E)** *Ndufa4l2* were correlated in the murine tumors. N=3 tumors.

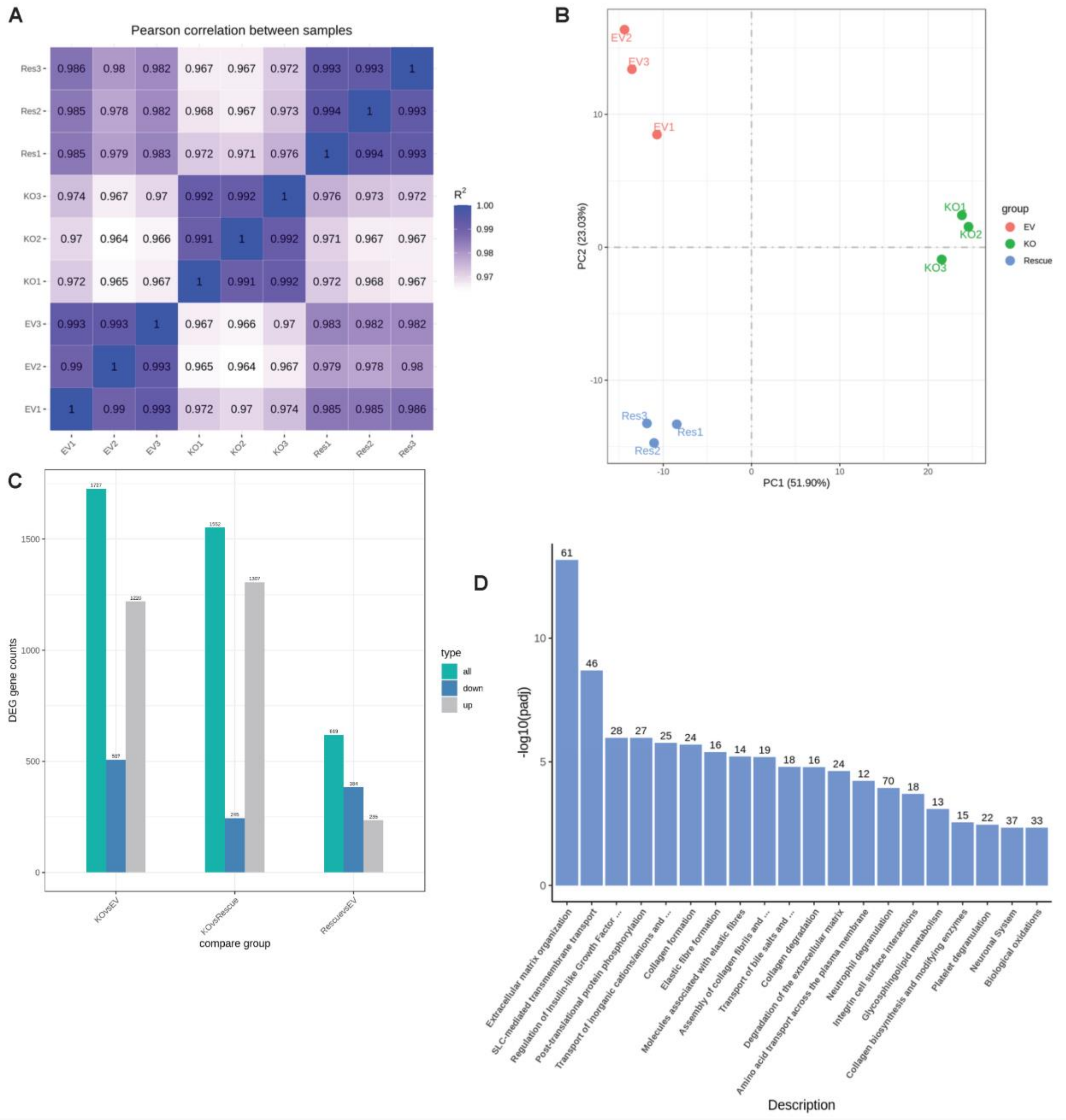


**Extended Data Figure 3. NDUFA4L2 rescue cells have similar expression profile to control cells.** Differential gene expression analysis from bulk RNA-sequencing gene counts from EV, NDUFA4L2 KO, and NDUFA4L2 rescue RENCA cells. **(A)** Pearson correlation heatmap between samples, **(B)** Gene cluster analysis, and **(C)** differential expression gene analysis comparing genes that are up or down in each group suggests that EV and rescue cells have similar gene expression profiles. **(D)** NDUFA4L2 KO compared to rescue in RENCA cells resulted in transcriptomic changes in metabolic and extracellular matrix pathways.

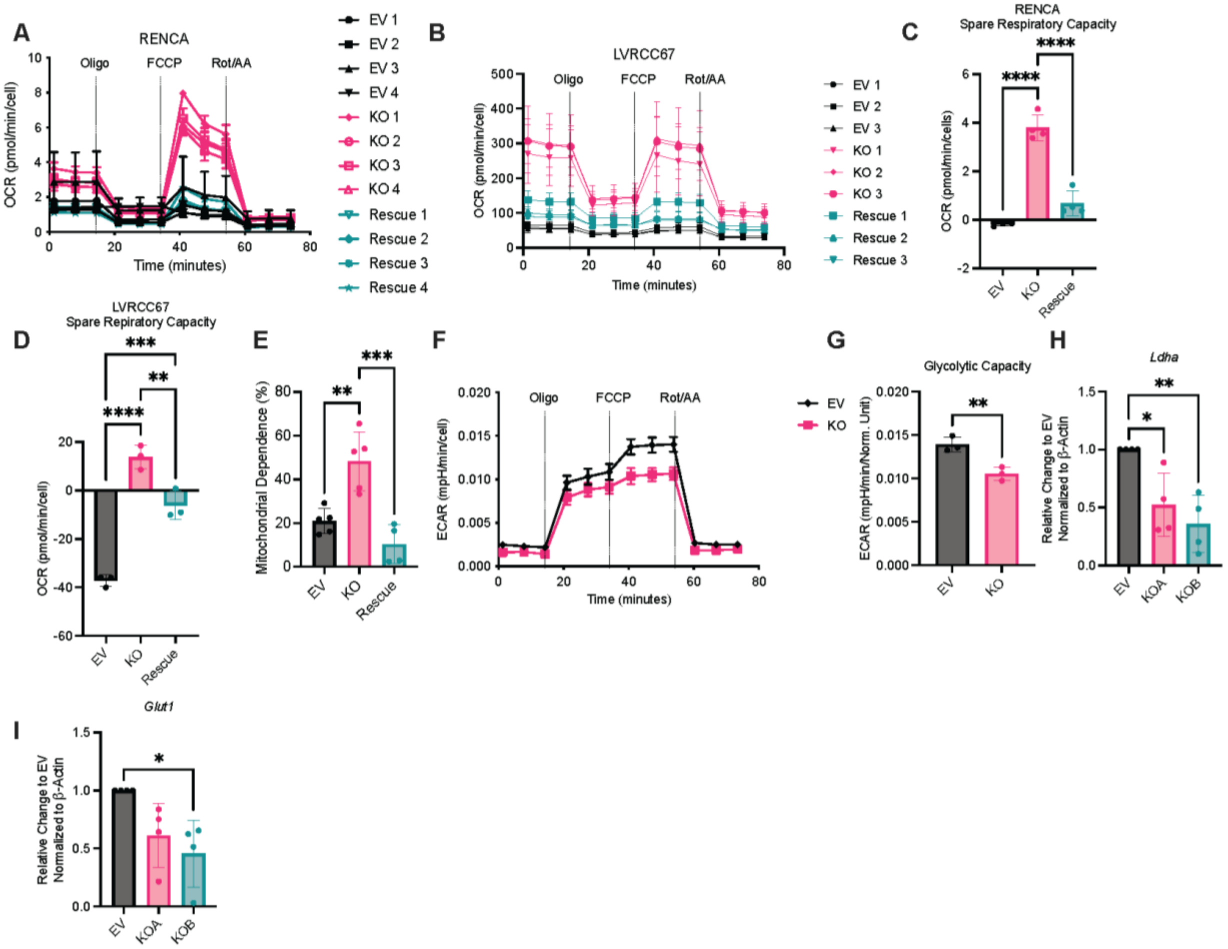


**Extended Data Figure 4. NDUFA4L2 expression correlates with glycolysis.** Extracellular flux mitostress test from **(A)** RENCA and **(B)** LVRCC67 EV, KO, and rescue cells. The graphs include 3-4 biological replicates, with 6 technical replicates/biological replicate each. OCR and ECAR measurements were normalized to total cell number in the well. In addition, spare respiratory capacity was measured from **(C)** RENCA and **(D)** LVRCC67 seahorse measurements. **(E)** SCENITH of EV, KO, and rescue RENCA cells treated with puromycin, 2DG, oligomycin individually and in combination. Protein translation was quantified by measuring puromycin incorporation in each condition and mitochondrial capacity (%) was calculated. Each dot represents a biological replicate combined from 2 individual experiments. **(F)** Seahorse bioanalyzer glyco-stress test from RENCA EV and KO RENCA cells. The graphs are representative of 3 or more individual experiments, with 3-4 biological replicates and 6 technical replicates each. **(G)** In addition, glycolytic capacity of EV and NDUFA4L2 KO RENCA cells was calculated from the seahorse measurements. Two different clonal RENCA NDUFA4L2 KO cell lines have glycolytic transcripts compared to empty vector including **(H)** *Ldha* and **(I)** *Glut1*. Gene expression was measured using qPCR and normalized to β-Actin. Fold Change was plotted by normalizing expression to the empty vector (EV). Each dot represents an individual experiment, with 3-6 technical replicates/experiment. Statistics were done using a one-way anova with multiple comparisons, *$p<0.05$, **$p<0.01$, ***$p<0.001$, ****$p<0.0001$.

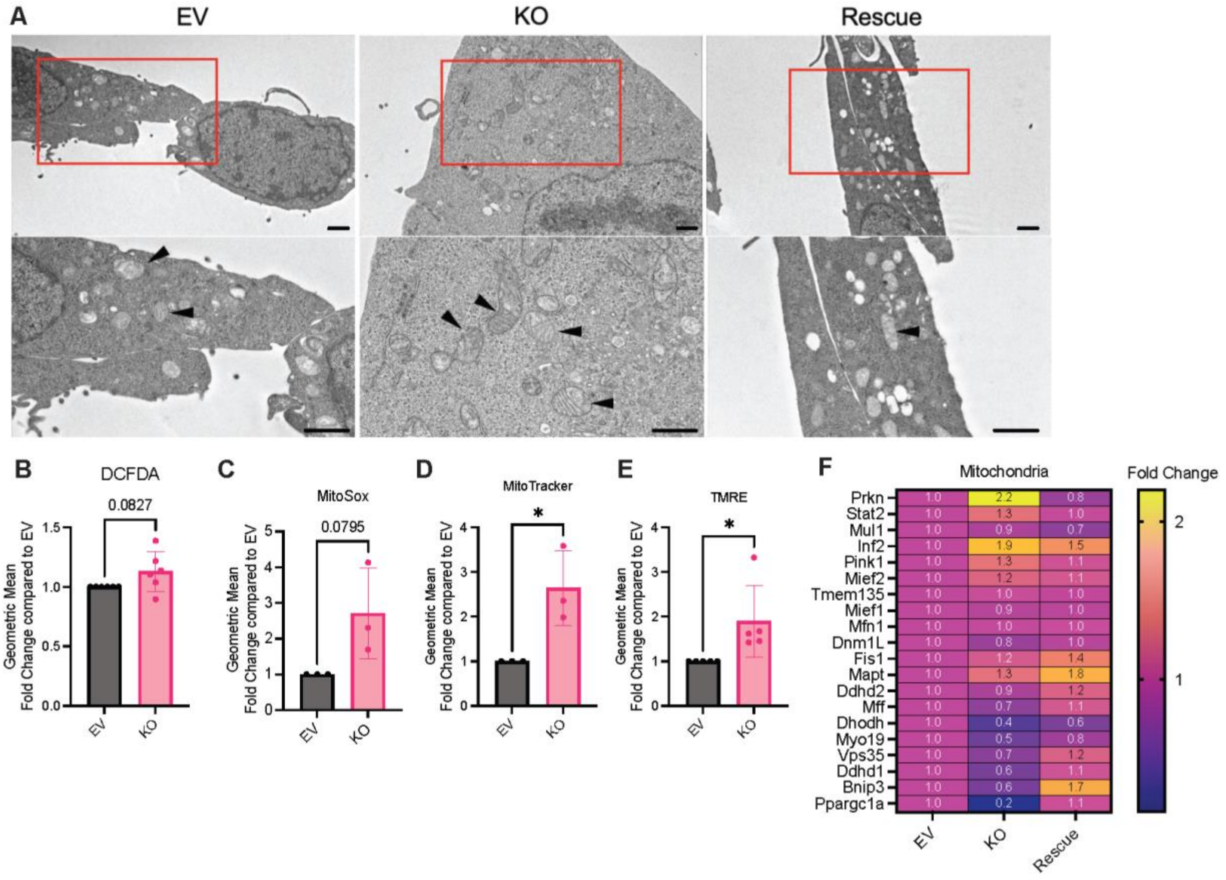


**Extended Data Figure 5. NDUFA4L2 regulates mitochondria. (A)** Mitochondria imaged using Transmission Electron Microscopy (TEM) in EV, KO, and rescue RENCA cells. Scale bars are 1μm. The red box indicates the area in which the zoomed in image was taken from. Geometric mean of **(B)** DCFDA (total cellular ROS), **(C)** MitoSox (mitochondrial ROS), **(D)** Mito tracker (mitochondrial mass), and **(E)** TMRE as a readout of mitochondria membrane potential in EV and NDUFA4L2 KO RENCA cells measured via flow cytometry. Each dot represents an individual experiment, with 6 replicates/experiment. (**F)** RNA-seq of RENCA EV, KO, and rescue cells revealed that NDUFA4L2 modulation influences mitochondria regulation genes. Three technical replicates were sequenced per cell line. Gene counts were normalized to the average of the EV gene counts. Statistics were done using a one-way anova with multiple comparisons, *$p<0.05$.

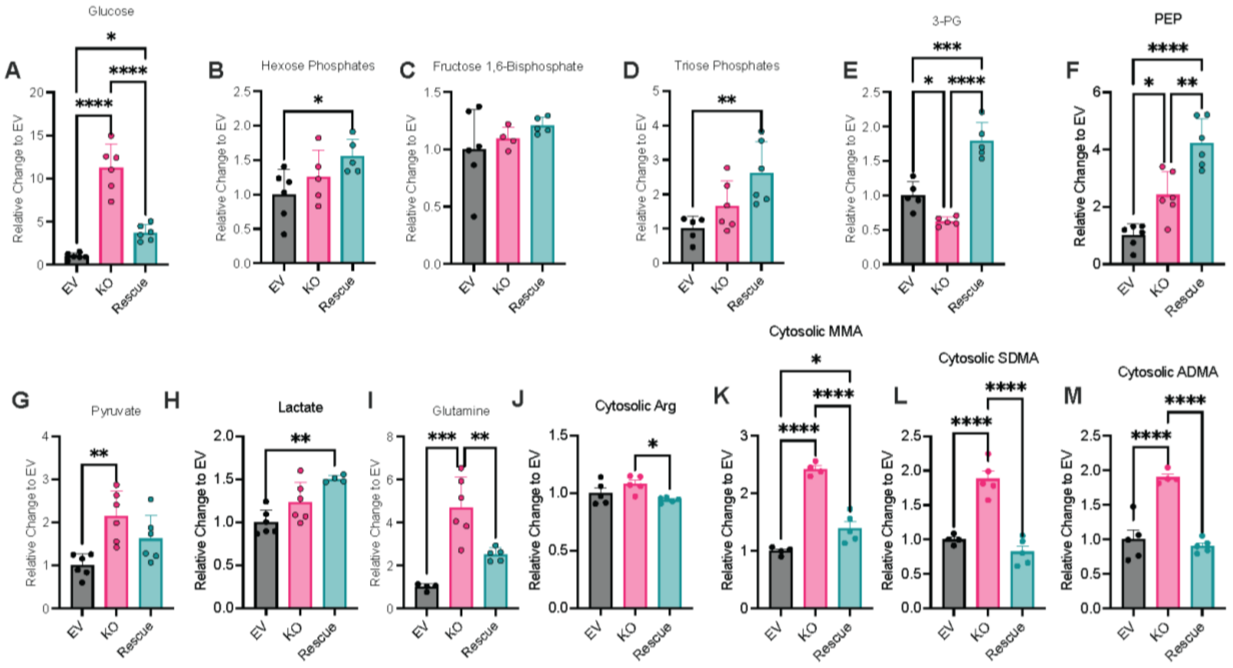


**Extended Data Figure 6. NDUFA4L2 expression results in metabolic and PTM changes.** LC-MS/MS was used to perform targeted metabolomics on EV, KO, and rescue RENCA cells. **(A)** glucose, **(B)** hexose phosphates, **(C)** fructose 1,6-bisphosphate, **(D)** triose phosphates, **(E)** 3-phosphoglycerate (3-PG), **(F)** phosphoenolpyruvate (PEP), **(G)** pyruvate, **(H)** lactate, and **(I)** glutamine were measured. All analytes were quantified using a $^{13}$C-1-Lactate internal standard and normalized to the protein in each respective sample's cell pellet. Relative change was calculated by normalizing to the average of the EV cells. N=5-6 replicates and is representative from 2 individual experiments. Arginine post-translational modifications were quantified in the cytosol of EV, NDUFA4L2 KO, and rescue RENCA cells. In the cytosol, **(J)** total arginine, **(K)** MMA, **(L)** SDMA, **(M)** and ADMA were measured. Relative change was calculated by normalizing to the average of the EV cells. N=4-5 replicates and is representative from 2 individual experiments. Statistics were done using a one-way anova with multiple comparisons, *$p<0.05$, **$p<0.01$, ***$p<0.001$, ****$p<0.0001$.

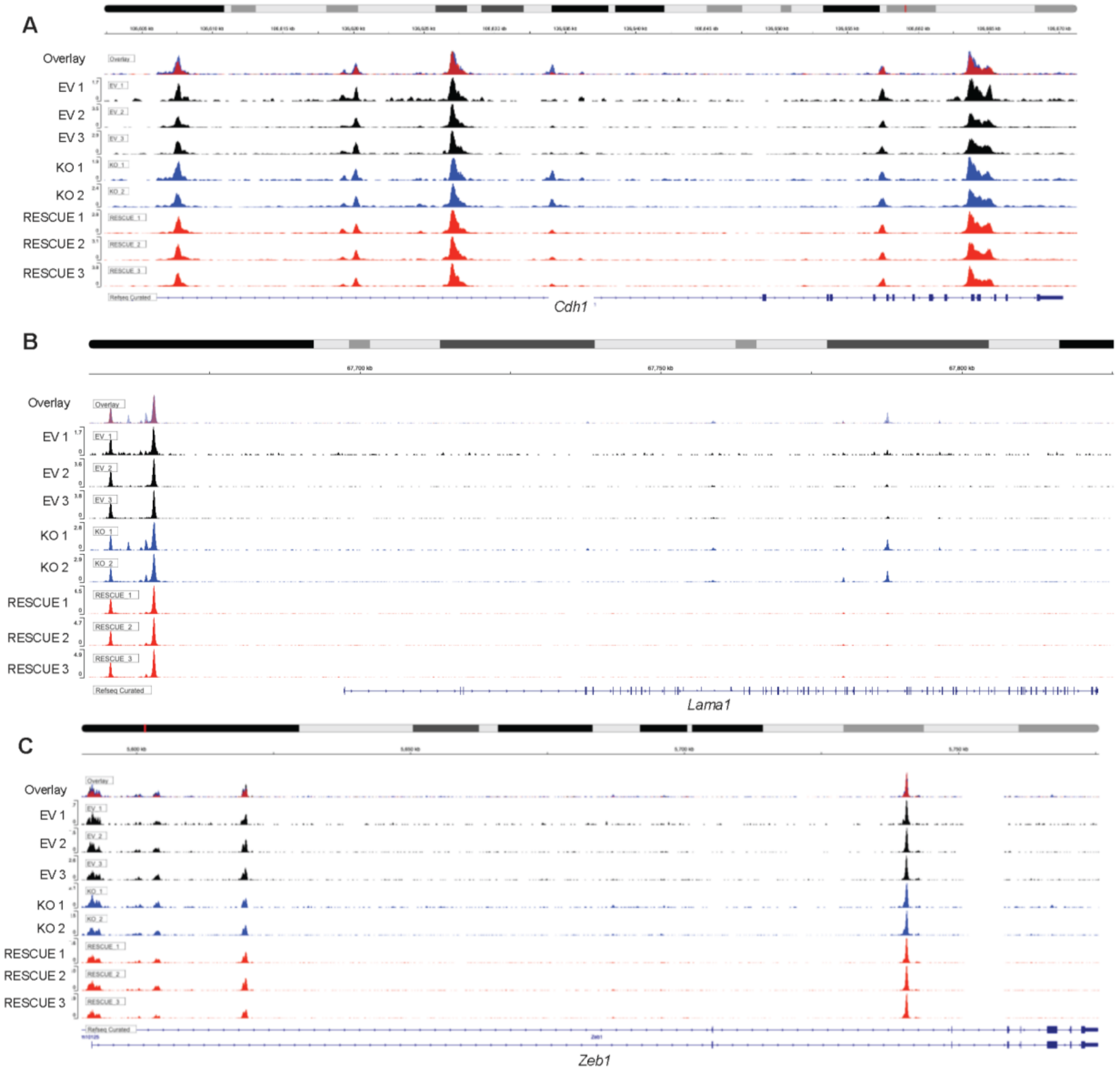


**Extended Data Figure 7. NDUFA4L2 regulates chromatin accessibility of EMT genes.** ATAC-sequencing plots for **(A)** *Cdh1*, **(B)** *Lama1*, and **(C)** *Zeb1*. Overlay is done with one representative biological replicate per group. Black is EV, blue in NDUFA4L2 KO, and red is rescue. 2-3 biological replicates were sequenced per group.

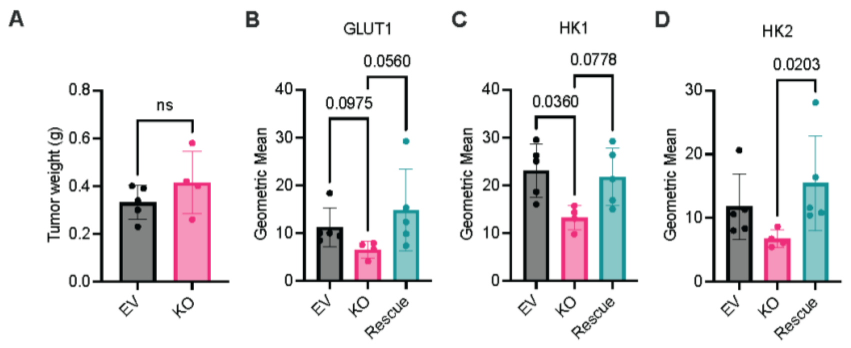


**Extended Data Figure 8. NDUFA4L2 KO cells retain decreased glycolytic enzymes ex vivo.** **(A)** Tumor weight of EV and NDUFA4L2 KO RENCA cells subcutaneously implanted into immunocompetent mice. Geometric mean of **(B)** *GLUT1*, **(C)** *HK1*, and **(D)** *HK2* on CD45 negative cells isolated from the orthotopic subrenal tumors in immunocompetent mice. Each dot represents a biological replicate, n = 5/group and is representative from two individual experiments. Statistics were done using a one-way anova and are individually listed for each comparison.

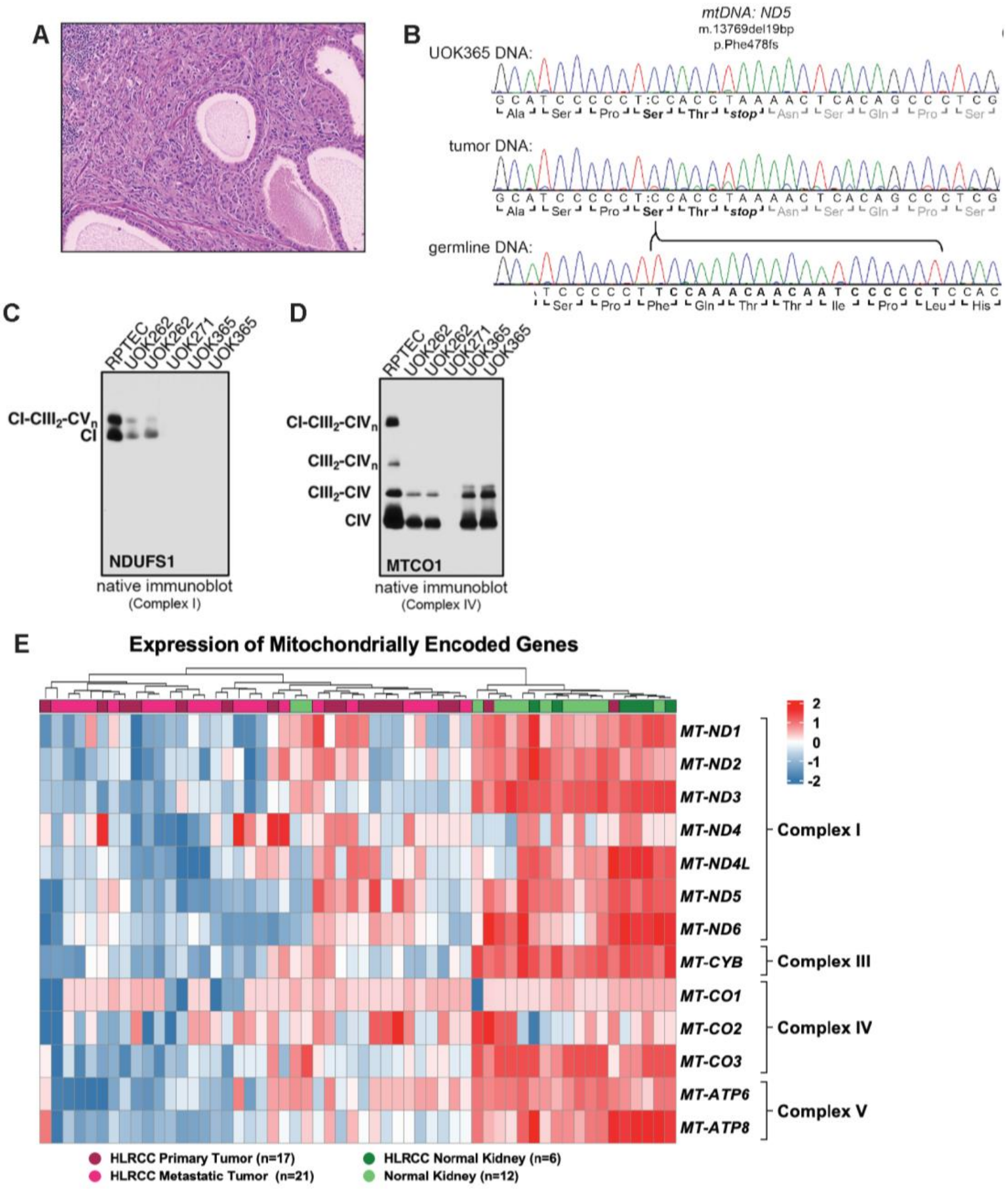


**Extended Data Figure 9. *FH*-deficient tumors have decreased ETC activity. (A)** Low magnification H&E staining of the HLRCC patient's tumor specimen revealed a solid/cystic histology. **(B)** $FH^{-/-}$ UOK365 cells were analyzed using both next generation and Sanger mtDNA sequencing and found to harbor a homoplasmic 19 base pair deletion in the mitochondrial encoded MT-ND5 gene, resulting in a frameshift mutation (p.Phe478fs) that truncates the MT-

ND5 protein product by 124 residues, or approx. 21% of the total protein length. The MT-ND5 mutation was also present in DNA obtained from the patient's tumor at a heteroplasmy level of 73.4% but was not present in the germline blood DNA. **(C)** Complex I assembly was evaluated by native immunoblot against the NDUFS1 subunit, which revealed diminished assembly of Complex I in UOK262 cells and undetectable Complex I levels in UOK271 and UOK365 cells. **(D)** Native immunoblot of Complex IV showing diminished assembly of CIV in UOK262 and undetectable CIV levels in UOK271 (MT-CO1 is a CIV copper and heme containing mtDNA encoded subunit). **(E)** Bulk-RNA sequencing was done on 17 primary HLRCC tumors, 12 HLRCC metastatic tumors, 6 HLRCC normal adjacent kidneys, and 12 normal kidneys from donors. Mitochondrially encoded genes for the electron transport chain were enriched in normal kidney compared with HLRCC tumors.

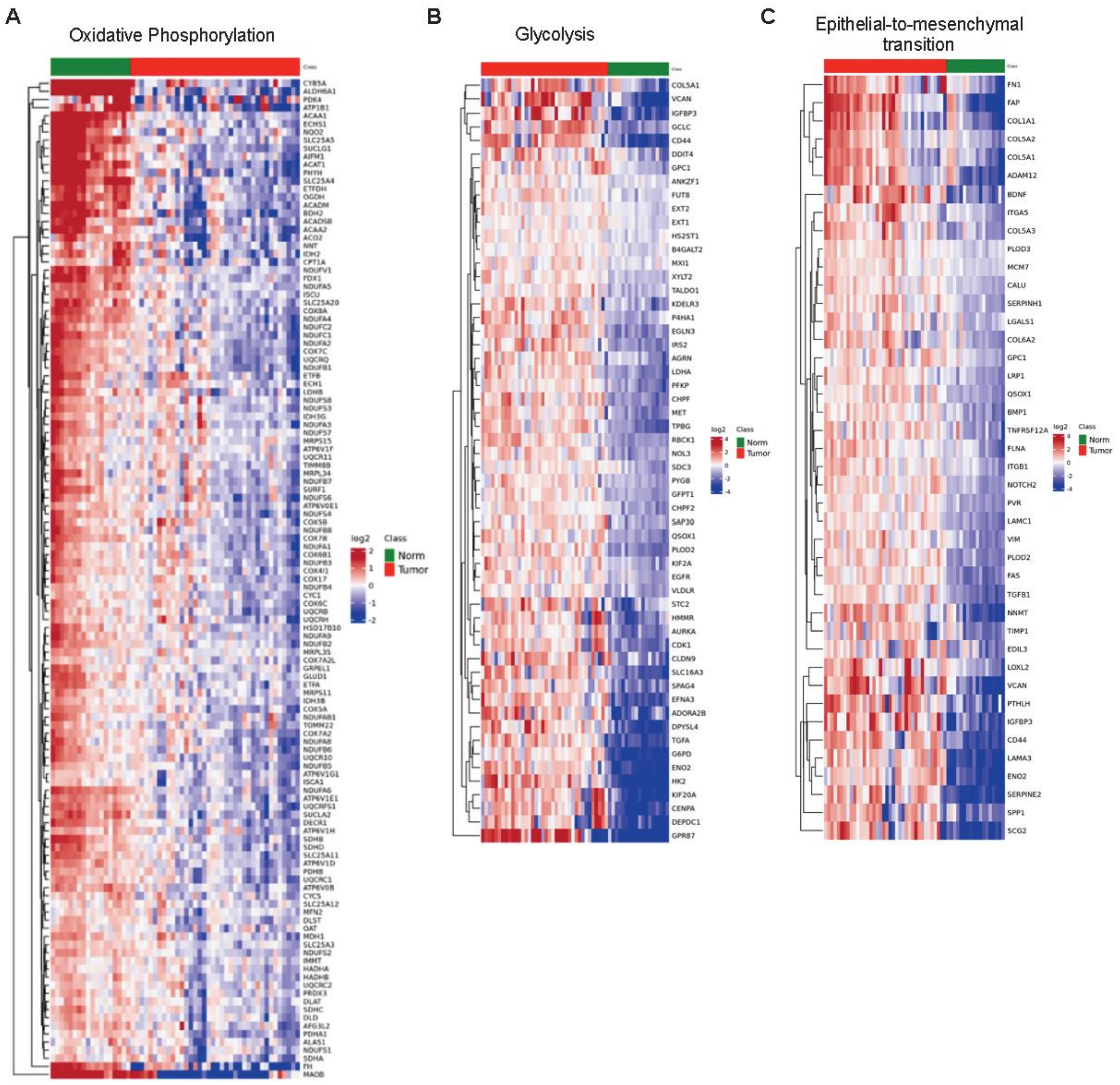


**Extended Data Figure 10. Loss of FH changes tumor cell metabolism and EMT.** Bulk-RNA sequencing was done on 17 primary HLRCC tumors, 12 HLRCC metastatic tumors, 6 HLRCC normal adjacent kidneys, and 12 normal kidneys from donors. Heatmaps from differentially expressed genes in **(A)** oxidative phosphorylation, **(B)** glycolysis, and **(C)** EMT pathways. In the bar at the top, green indicates normal kidney samples and red indicates HLRCC tumor samples.

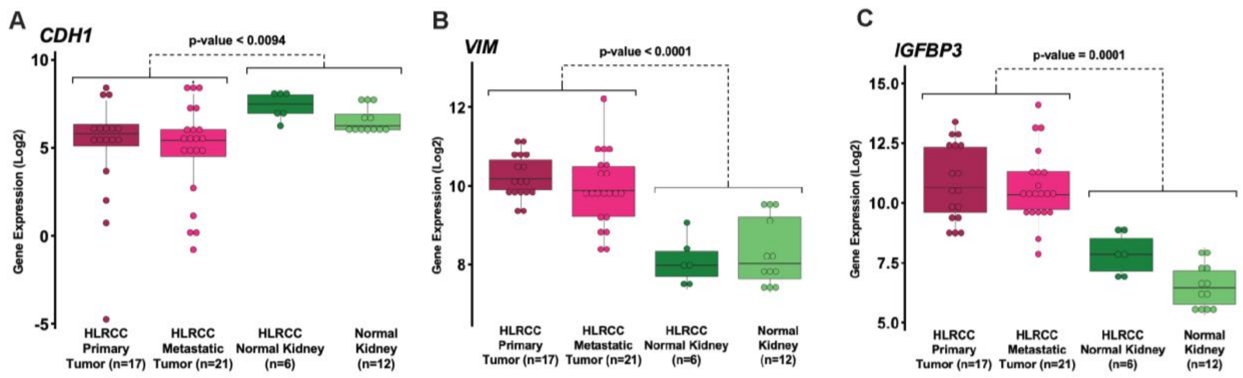


**Extended Data Figure 11. Loss of FH increases EMT.** Bulk-RNA sequencing was done on 17 primary HLRCC tumors, 12 HLRCC metastatic tumors, 6 HLRCC normal adjacent kidneys, and 12 normal kidneys from donors. Gene expression of **(A)** *CDH1*, **(B)** *VIM*, and **(C)** *IGFBP3* in each group. Statistic were doing using a one-way anova.

| Target | Method | Concentration | Catalog Number |
|---|---|---|---|
| **E-Cadherin** | Western | 1:1,000 | Cell Signaling 3195 |
| **β-Actin** | Western | 1:2,000 | Sigma A2066 |
| **Phalloidin 546** | Immunofluorescence | 1:500 | Invitrogen A22283 |
| **Hoechst 33342** | Immunofluorescence | 1:10,000 of 20mM Stock | Invitrogen 62249 |
| **E-Cadherin** | IHC | 1:500 | Cell Signaling 3195 |
| **Slug** | IHC | 1:500 | Cell Signaling 9585 |
| **CM-H2DCFDA** | Flow Cytometry | Reconstitute in 43uL DMSO, use 12.5uL/5mL PBS | Thermofisher D399 |
| **MitoSox** | Flow Cytometry | Reconstitute in 13uL DMSO, use 5uL/5mL PBS | Thermofisher M36008 |
| **Mitotracker Green FM** | Flow Cytometry | Reconstitute in 70uL DMSO (for 1mM stock), use 1uL/5mL PBS | Thermofisher M7514 |
| **TMRE** | Flow Cytometry | From 15mM stock, dilute 3uL in 2.25mL PBs, then dilute 50uL in 4.95mL PBS | LifeTech T-669 |
| **CD45 BV510** | Flow Cytometry | 1:1,600 | BioLegend 103138 |

| | | | |
|---|---|---|---|
| **Glut1 AF647** | Flow Cytometry | 1:500 | Abcam ab195020 |
| **HK1 AF647** | Flow Cytometry | 1:100 | Abcam ab197864 |
| **HK2 AF647** | Flow Cytometry | 1:200 | Abcam ab237314 |
| **Ef780 Viability Dye** | Flow Cytometry | 1:3,000 | BioLegend 65-0865-14 |

**Table S1. Antibodies used in each study.**

| Species | Q1 (m/z) | Q3 (m/z) | RT (min) |
|---|---|---|---|
| **Glucose** | 179 | 89 | 6.9 |
| **Hexose Phosphates** | 259 | 97 | 13.4 |
| **Fructose 1,6-Bisphosphate** | 339 | 97 | 13.8 |
| **Triose Phosphates** | 169 | 79 | 13 |
| **3-Phosphoglycerate** | 185 | 97 | 13.4 |
| **Phosphoenolpyruvate** | 167 | 79 | 13.3 |
| **Pyruvate** | 87 | 43 | 3.0 |
| **Lactate** | 89 | 43 | 6.0 |
| **$^{13}$C-1-Lactate (Internal Standard)** | 90 | 43 | 6.0 |
| **a-Ketoglutarate** | 145 | 101 | 10.3 |
| **Succinate** | 117 | 73 | 10.9 |
| **Glutamine** | 145 | 127 | 9.5 |

**Table S2. Analytes measured with targeted metabolomics**.

| Species | Q1 (m/z) | Q3 (m/z) |
|---|---|---|
| **Arg** | 175 | 70 |
| **$^{13}C_6{}^{15}N_4$-Arg [INTERNAL STANDARD]** | 185 | 75 |
| **MMA** | 189 | 70 |
| **ADMA** | 203 | 70 |
| **SDMA** | 203 | 172 |
| **Leu** | 132 | 86 |
| **$^{13}C_6{}^{15}N$-Leu [INTERNAL STANDARD]** | 139 | 93 |

**Table S3. Post-translational modifications measured with LC-MS/MS.**

| Gene Target | Forward Primer | Reverse Primer |
|---|---|---|
| **NDUFA4L2** | GTCTAGCAGCATCAGTCCAGC | CCACACCCAGTCAGGAGGAG |
| **CDH1** | CTGCTGCTCCTACTGTTTCTAC | TCTTCTTCTCCACCTCCTTCT |
| **CDH2** | AGTGGCAGGTAGCTGTAAAC | TGGCAAGTTGTCTAGGGAATAC |
| **β-Actin** | TCAAGATCATTGCTCCTCCTGAGC | TACTCCTGCTTGCTGATCCACATC |
| **GLUT1** | TCAACGAGCATCTTCGAGAAGGCA | TCGTCCAGCTCGCTCTACAACAAA |
| **LDHA** | ACAAACTCAAGGGCGAGATG | GGAGTTCGCAGTTACACAGTAG |

**Table S4. qRT-PCR Primer Sequences.**
